# One-step glycan photoaffinity probe synthesis enables identification of human milk oligosaccharide-interacting microbiota

**DOI:** 10.64898/2026.10.31.763344

**Authors:** Yuge Chun, Jack T. Leonard, Matthew E. Griffin

## Abstract

Glycans are found on the surfaces of all living cells and engage in cell signaling through the binding of various molecular targets. However, chemical probes to detect and identify carbohydrate interactions remain difficult to produce, due in large part to the challenges of late-stage, site-specific functionalization of oligosaccharides. Here, we report the synthesis and application of a minimal bifunctional diazirine/alkyne linker utilizing an *N,O*-dialkylhydroxylamine (NODHA) moiety that can readily be installed on the anomeric position of unprotected sugars. Our linker functionalizes glycans with complete β-stereoselectivity, appends less mass than a single hexose unit, and maintains the closed-ring structure of the reducing end sugar. With this molecule, we generate glycan photoaffinity probes in a single step using unprotected human milk oligosaccharides (HMOs). We then validate binding selectivity of our compounds using lectins with known affinity towards our sugars. Finally, we demonstrate the utility of these glycan photoaffinity probes by performing photocrosslinking on a complex microbiota community and use cell sorting paired with metagenomic sequencing to identify potential new HMO-interacting microbes. Together, our work provides a one-step route to synthesize photoaffinity probes from unprotected, reducing glycans for interrogating carbohydrate interactions at the host-microbiota interface.

## Introduction

Carbohydrates are ubiquitous molecules on virtually all cell surfaces that can elicit cell-cell communication by engaging both soluble proteins and transmembrane receptors.^1^ These interactions regulate cellular, tissue, and organ physiology and contribute to numerous essential processes including immune cell development,^2^ vascular homeostasis,^3^ and barrier integrity.^4^ Glycans also mediate interactions between microbes and host organisms, enabling toxin binding,^5^ antibody recognition,^6^ and commensal colonization.^7^ Moreover, certain soluble glycans from both host and microbial sources are recognized as potent signaling molecules. For example, fragments of bacterial glycopeptides^8-10^ and precursors of microbial sugar biosynthesis^11^ can activate host innate immune responses through highly conserved mammalian receptors. In addition, host cells can produce carbohydrates that are either secreted or enzymatically liberated by microbes to foster cross-feeding and promote selective proliferation.^12-14^ Therefore, the identification of carbohydrate interactions is crucial to understand molecular mechanisms of cell signaling both within and between kingdoms of life.

Compared to interactions between proteins, carbohydrate binding is generally a lower affinity phenomenon depending on glycan size and composition, with apparent dissociation constants for oligosaccharides varying widely across the millimolar to mid-micromolar regime.^15^ These interactions are thus more transient and dynamic, which impedes their detection through traditional purification methodologies that rely on high-affinity binding. To overcome this challenge, elegant solutions such as glycodendrimers and glycopolymers have been developed.^16^ Like natural repeating patterns in polysaccharides, these methods exploit principles of artificial multivalency to increase the local concentration of lower affinity glycan ligands and facilitate target binding and detection. Despite their utility, these methods require both the synthesis or isolation of pure oligosaccharides and their subsequent attachment to an additional synthetic scaffold, which may not recapitulate the natural structure of smaller glycan fragments used as signaling molecules.

Alternatively, photocrosslinking has emerged as a powerful tool to capture transient interactions including carbohydrate binding.^17^ This approach most commonly utilizes bifunctional linkers that contain a photoactivable group like a diazirine to crosslink with target proteins upon UV irradiation as well as another moiety such as an alkyne handle to facilitate downstream labeling or purification of the covalently bound target.

Photocrosslinking probes have been highly successful at identifying glycan-binding targets.^18^ Nevertheless, glycan derivatives are quite difficult to synthesize de novo due to the numerous protection, glycosylation, and deprotection steps required to assemble a complex oligosaccharide and append the functional moieties with regio- and diastereoselectivity.^19, 20^ Bifunctional photoaffinity handles often cannot tolerate these transformations and require late-stage modification. Therefore, methods to directly modify unprotected synthetic sugars or naturally derived glycans would greatly accelerate our ability to identify carbohydrate binding partners and decipher their biological functions.

Reductive amination has been widely used to functionalize unprotected sugars (**Figure 1A**). However, this approach permanently opens the ring of the reducing end monosaccharide, which may alter binding affinity towards its receptors. In response, two main strategies have been developed to form closed-ring glycoside probes and preserve the native structures of the glycan units. First, glycoconjugates can be produced using a two-step approach that first forms an anomeric hemiaminal followed by amidation (**Figure 1B**).^21^ The Townsend, Parker, and Huang labs employed this method to produce photoaffinity probes of human milk oligosaccharides (HMOs), which are prevalent components of human breast milk that can be isolated as unmodified sugars.^22, 23^ Although powerful, the two-step sequence exhibited moderate overall yields of 10% to 37% across substrates when employing classical Kochetkov amination conditions to produce the glycosylamine intermediate. These yields were improved to 44% to 53% using optimized microwave conditions along with a slight excess of the glycosylamines for the amide coupling, which may be limiting for more precious oligosaccharide starting materials.

**Figure 1.**
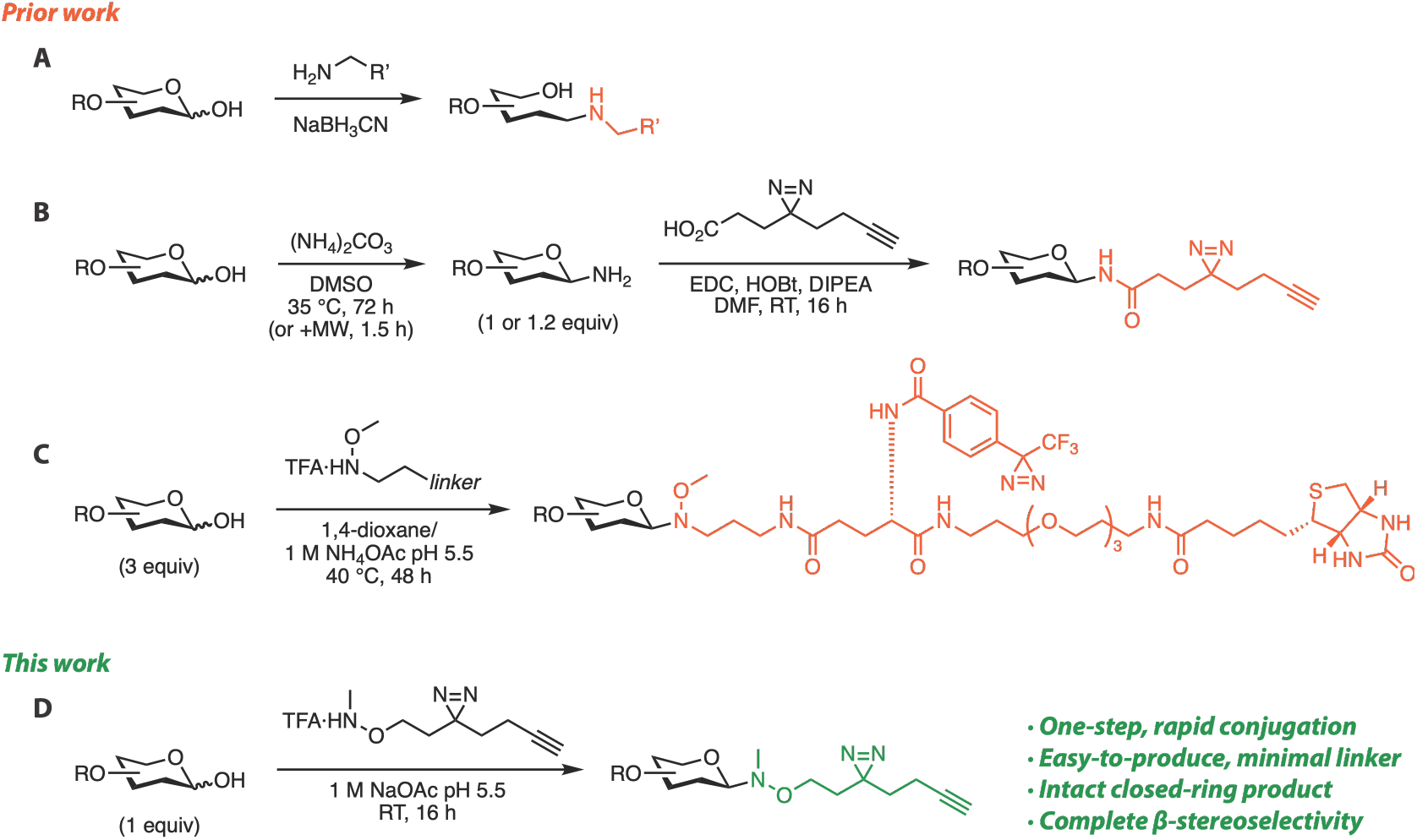
Prior and current methods to synthesize glycan probes using unprotected sugars. **(A)** Reductive amination of reducing sugars. **(B)** Two-step Kochetkev amination and amidation of reducing sugars to produce photoaffinity glycan probes, MW = microwave. **(C)** One-step *N,O-*dialkylhydroxyamine (NODHA) conjugation of reducing sugars to produce photocrosslinking and biotinylated glycan probes. **(D)** One-step NODHA conjugation using a minimal bifunctional photoaffinity linker.

Second, *N,O*-dialkylhydroxylamine (NODHA) linkers have been developed to generate diverse glycan probes in a single step.^24-30^ These reactions involve the formation of an oxyiminium intermediate by reaction of the NODHA nitrogen nucleophile with the open-ring carbonyl form of reducing sugars (**Scheme S1**). The electrophile is then trapped by the former ring oxygen in a 6-exo-trig cyclization process to form a stable, closed-ring *N*-glycoside. This approach has been broadly successful to develop many glycoscience tools, including carbohydrate microarrays,^24-27^ glycopolymers,^28^ and “clickable” probes for further functionalization.^27, 29^ In addition, biotin-containing photocrosslinkers have also been installed onto unprotected glycans using NODHA chemistry, which enabled streptavidin-mediated detection of crosslinked lectins (**Figure 1C**).^30^ Despite their functionality, these multifunctional probes are often bulky, with molecular weights equal to or even larger than the glycan moiety. Large chemical handles may in turn alter interactions or molecular distributions in biological settings. Therefore, we aimed to produce a minimal photoaffinity tag that could be installed in a single step onto complex, unprotected oligosaccharides. We envisioned that this late-stage strategy using a minimal tag would not only circumvent the difficulties of de novo glycoconjugate synthesis but also avoid disrupting the closed ring structure of the reducing monosaccharide or altering its binding affinity.

Here, we describe the synthesis of a new class of bifunctional diazirine/alkyne linkers based on the NODHA scaffold for the one-step production of photoaffinity glycan probes (**Figure 1D**). We first identified functionalization conditions that are tolerated by the reactive diazirine moiety. We then optimized the conjugation reaction using an unprotected disaccharide starting material, focusing only on conditions where the more precious glycan acted as the limiting reactant. Next, we explored several structural variants of our linker to identify factors that may alter the yield of the conjugation reaction or efficiency of photocrosslinking. We produced several glycan photoaffinity probes based on dietary HMOs and observed selective, proof-of-principle photocrosslinking of glycan-binding proteins. We used our probes to demonstrate labeling of specific microbes within a complex microbiota in a structure-dependent manner. Finally, we show that this approach, combined with targeted cell sorting and metagenomic sequencing, can enable the identification of glycan-binding microbiota. Together, our work reveals an expedient method to produce oligosaccharide photoaffinity probes that can be readily adapted to more structurally complex reducing sugars.

## Results and Discussion

To synthesize our initial linker *N*-Me-*O*-DiazAlk linker **1** (**Scheme 1**), *N-*Boc*-N*- methylhydroxylamine was conjugated to 3-(but-3-yn-1-yl)-3-(2-iodoethyl)-3H-diazirine using DBU-mediated S_N_2 conditions. Deprotection of the Boc group by 5% TFA yielded **1** in an overall yield of 70% over two steps. Importantly, the two-step sequence to access **1** uses facile chemistry with commercially available starting materials, making this compound easily accessible to the broader community.

**Scheme 1.**
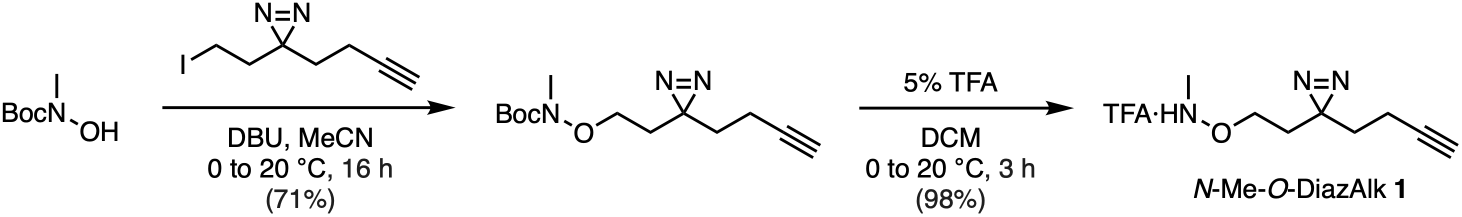
Synthesis of diazirine/alkyne bifunctional linker *N*-Me-*O*-DiazAlk **1**.

Previous reports of conjugation reactions using NODHA linkers have employed various acidic conditions with elevated temperatures for 2–72 hours.^24-30^ Initially, we followed previously reported conditions (DMSO/AcOH, 7:3, v/v)^26^ to attach **1** onto lactose as a model sugar substrate with 0.5 to 2.0 equivalents of the linker (**Table S1**). Crude NMR analysis suggested that these conditions resulted in only moderate conversion values for target products with an apparent loss of **1**. Therefore, we postulated that the diazirine moiety may be unstable under these prolonged incubations. We then systematically tested the stability of our linker under three commonly used acidic conditions at different temperatures. We monitored the progress of these reactions by ^1^H NMR spectroscopy using dimethylsulfone (DMS) as an internal standard to quantify remaining linker in the crude reaction mixture (**Table 1**). Rapid decomposition of **1** was observed in DMSO containing 30% AcOH even at room temperature (Entry 1; **Table 1**). Conversely, aqueous conditions containing 1 M sodium acetate (NaOAc) buffer at pH 5.0 or a 1:1 mixture of 1,4-dioxane and 1 M ammonium acetate (NH_4_OAc) buffer at pH 5.5 showed little to no appreciable degradation of **1** after 16 h at room temperature (Entries 3 and 5; **Table 1**). However, elevated temperatures in both buffer systems led to progressively lower recoveries of **1** (Entries 4, 5, 7, and 8; **Table 1**). Together, these results suggest **1** is sufficiently stable at room temperature under mildly acidic conditions for use in our conjugation reactions.

**Table 1.** Stability profiling of diazirine linker *N*-Me-*O*-DiazAlk **1**.

| Entry | Solvent | Temperature (°C) | **1** remaining (%)<sup>[1]</sup> |
|---|---|---|---|
| 1 | DMSO/AcOH (7:3 v/v) | 25 | 0 |
| 2 |  | 40 | 0 |
| 3 | 1 M NaOAc pH 5.0 | 25 | 96 |
| 4 |  | 40 | 79 |
| 5 |  | 65 | 0 |
| 6 | 1,4-Dioxane and<br>1 M NH<sub>4</sub>OAc pH 5.5 (1:1 v/v) | 25 | 91 |
| 7 |  | 40 | 79 |
| 8 |  | 65 | 28 |

1. The amount of remaining starting material was calculated by <sup>1</sup>H NMR analysis of crude reaction mixtures using dimethylsulfone (DMS) as an internal standard (see **Table S2**).

After identifying conditions in which **1** was stable, we proceeded to further optimize the conjugation reaction between **1** and reducing sugars. Using the disaccharide lactose as a model substrate, we performed reactions using both acetate buffers with varying temperature, linker stoichiometry, and additives (**Table 2**). We used lactose as the limiting reactant in these reactions to mimic conditions in which precious oligosaccharides would be used in the future. Conversion values were determined by comparing the ^1^H NMR integration values for signals corresponding to the anomeric protons of the product and the remaining starting material in crude reaction mixtures. Treatment of lactose with two equivalents of **1** in 1 M NaOAc pH 5.0 at room temperature afforded the desired glycoconjugate **Lac-1** in 46% conversion after 16 hours (Entry 1, **Table 2**). Increasing the amount of **1** to four equivalents resulted in an improved conversion of 65% (Entry 2, **Table 2**). Higher amounts of **1** proved impractical due to solubility issues in the reaction buffer. Increasing reaction temperature to 40 °C provided only moderate enhancement in conversion to 72% (Entry 4, **Table 2**). When the pH of the initial buffer system was decreased to pH 4.0, we observed poorer yields at room temperature and degradation with gentle heating (Entries 6 and 7, **Table 2**), indicating the pH dependence of NODHA glycoside formation. Using a 1:1 mixture of 1,4-dioxane and 1 M NH_4_OAc pH 5.5 led to a mixture of alpha and beta anomers (Entries 8 and 9, **Table 2**).

**Table 2.**
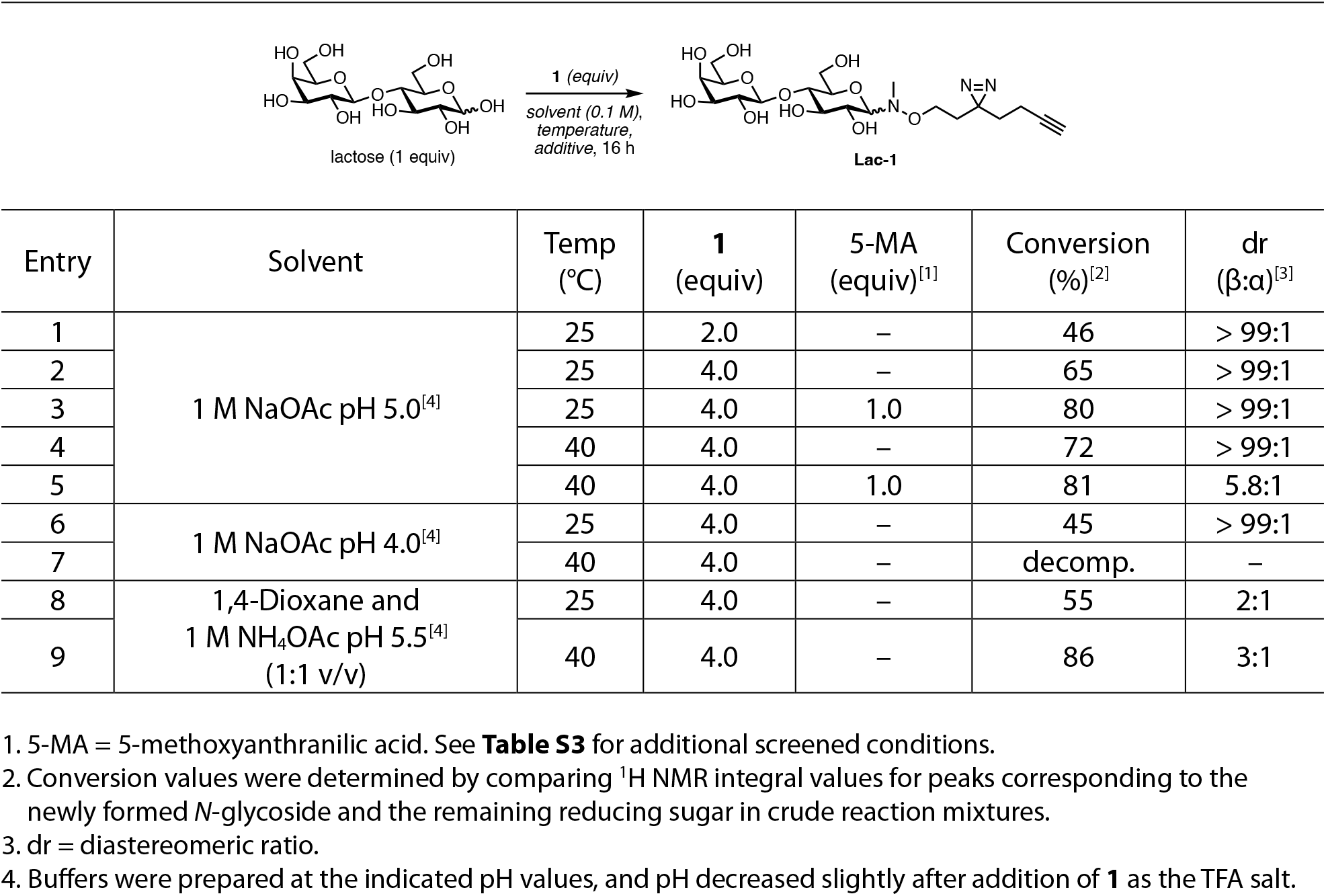
Optimization of conjugation conditions of lactose with **1**.

| Entry | Solvent | Temp<br>(°C) | **1**<br>(equiv) | 5-MA<br>(equiv)<sup>[1]</sup> | Conversion<br>(%)<sup>[2]</sup> | dr<br>(β:α)<sup>[3]</sup> |
|---|---|---|---|---|---|---|
| 1 | 1 M NaOAc pH 5.0<sup>[4]</sup> | 25 | 2.0 | – | 46 | > 99:1 |
| 2 |  | 25 | 4.0 | – | 65 | > 99:1 |
| 3 |  | 25 | 4.0 | 1.0 | 80 | > 99:1 |
| 4 |  | 40 | 4.0 | – | 72 | > 99:1 |
| 5 |  | 40 | 4.0 | 1.0 | 81 | 5.8:1 |
| 6 | 1 M NaOAc pH 4.0<sup>[4]</sup> | 25 | 4.0 | – | 45 | > 99:1 |
| 7 |  | 40 | 4.0 | – | decomp. | – |
| 8 | 1,4-Dioxane and<br>1 M NH<sub>4</sub>OAc pH 5.5<sup>[4]</sup><br>(1:1 v/v) | 25 | 4.0 | – | 55 | 2:1 |
| 9 |  | 40 | 4.0 | – | 86 | 3:1 |

1. 5-MA = 5-methoxyanthranilic acid. See **Table S3** for additional screened conditions.
2. Conversion values were determined by comparing <sup>1</sup>H NMR integral values for peaks corresponding to the newly formed *N*-glycoside and the remaining reducing sugar in crude reaction mixtures.
3. dr = diastereomeric ratio.
4. Buffers were prepared at the indicated pH values, and pH decreased slightly after addition of **1** as the TFA salt.

Finally, prior work had found that the addition of 5-methoxyanthranilic acid (5-MA) could improve yields of NODHA conjugation reactions.^31, 32^ We were able to reproduce this improvement in our hands as well using one equivalent of 5-MA; however, use of this reagent at 40 °C led to a mixture of diastereomers (Entries 3 and 5, **Table 2**). Therefore, we moved forward with conditions in Entry 3 to maximize yield, minimize degradation of **1**, and maintain β-stereoselectivity.

With optimization complete, we began to scale up the production of **Lac-1** for photocrosslinking experiments. To purify our highly hydrophilic products, we optimized a two-step organic and solid-phase extraction protocol to recover excess unreacted **1** and remove unreacted glycan from our product, respectively. Unfortunately, separation of the unreacted starting material, conjugated product, and 5-MA proved quite laborious and low-yielding in our hands even after extensive optimization of the extraction and separation protocols. Therefore, we chose to conduct all subsequent reactions and purifications using conditions reported in Entry 2 without 5-MA.

With our initial **Lac-1** probe in hand, we validated its ability to photocrosslink glycan-binding proteins using the plant lectins, soybean agglutinin (SBA) and *Lotus tetragonolobus* lectin (LTL) (**Figure 2a**). The preferred glycan recognition motif for SBA is terminal GalNAc; however, lactosylamine resin can be used to purify SBA due to residual binding affinity.^33^ Similarly, the canonical recognition of LTL has been described as the Lewis^X^ structure, which contains the lactosyl group.^34^ Therefore, these lectins function as stringent screens for binding and photocrosslinking. Lectins (1 mg mL^-1^) were incubated with **Lac-1** at varying concentrations for 20 minutes at 0 °C followed by UV irradiation for 10 minutes at 0 ºC. Labeled lectins were then reacted with biotin-PEG_3-_azide using copper-catalyzed azide-alkyne cycloaddition (CuAAC) conditions, resolved by SDS-PAGE, and detected by streptavidin blotting. **Lac-1** showed dose-dependent photocrosslinking to both lectins, and signals were dependent on UV irradiation (**Figure 2b**). In addition, photocrosslinking signals on both lectins decreased with increasing concentrations of unmodified lactose (**Figure 2c**), demonstrating that **Lac-1** and lactose compete for the same binding site.

**Figure 2.**
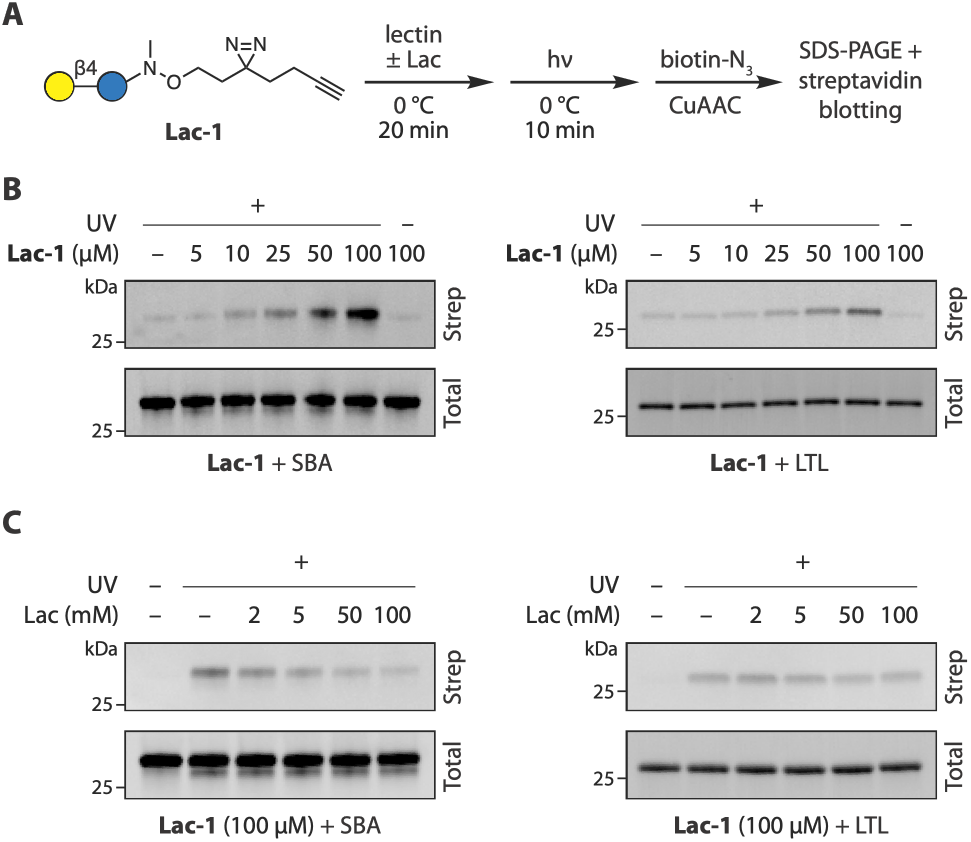
Photocrosslinking of **Lac-1** to glycan binding proteins, soybean agglutinin (SBA) and *Lotus tetragonolobus* lectin (LTL). **(A)** Workflow of photocrosslinking experiments. **(B)** Photocrosslinking of **Lac-1** to SBA or LTL occurs in a UV- and dose-dependent manner. **(C)** Presence of excess unmodified lactose can outcompete crosslinking of **Lac-1** to SBA or LTL.

In addition to a linear minimalist linker, prior reports by the Parker lab have shown that “split” minimalist linkers in which the diazirine and alkyne groups are separated may improve photocrosslinking and subsequent target identification by mass spectrometry.^35^ Therefore, we designed two additional photoaffinity linkers *N*-Alk-*O*-Diaz **2** and *N*-Diaz-*O*-Alk **3** bearing different connectivities of the two functional groups on the *N*- and *O*-substituents of the hydroxylamine group. Boc-protected versions of these linkers were produced via sequential S_N_2 substitution reactions to install each arm, furnishing the desired compounds in 49% and 40% yield over two steps, respectively (**Schemes 2a,b**). These linkers were treated with TFA and then immediately conjugated to lactose using the optimized conditions to produce **Lac-2** and **Lac-3** in 14% and 13% yield, respectively (**Scheme 2c**). Reaction yields for the branched linkers **2** and **3** were notably lower than the linear linker **1** and reflected lower conversions, which may be due to increased steric bulk near the nitrogen nucleophile.

**Scheme 2.**
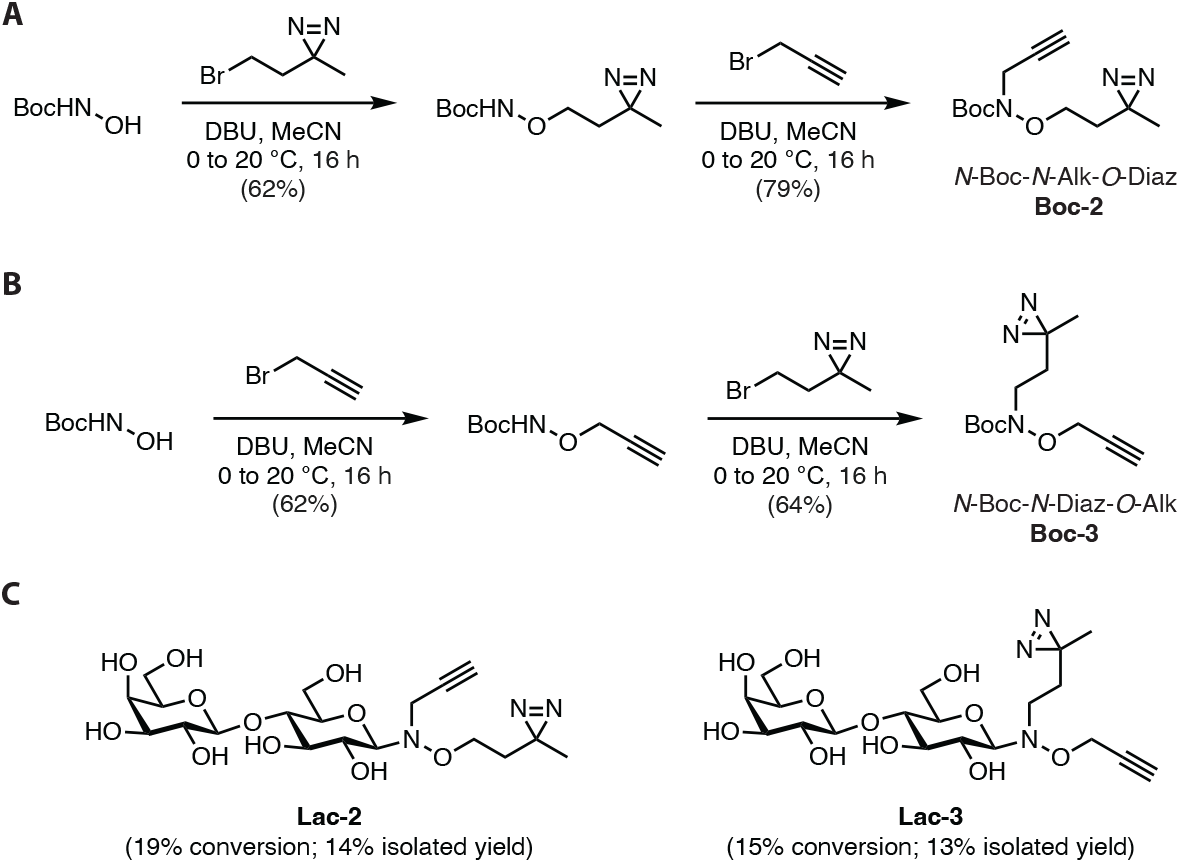
**(A, B)** Synthesis of Boc-protected versions of split diazirine/alkyne bifunctional linkers *N*-Alk-*O*-Diaz **2** and *N*-Diaz-*O*-Alk **3. (C)** Structures and yields of **Lac-2** and **Lac-3**.

Despite lower yields, we were able to generate enough of each lactose probe to compare their ability to crosslink with a target protein. **Lac-1, Lac-2**, and **Lac-3** were each subjected to the previous photocrosslinking workflow using LTL (**Figure 3a**). Quantification of blot intensities revealed that **Lac-3** crosslinked to LTL with approximately 2-fold higher efficiency compared to that of **Lac-1** or **Lac-2** (**Figure 3b**). This difference in activity may be due to the shorter distance between the latent carbene formed by UV irradiation and the glycan (4 atoms for **Lac-3** and 5 atoms for **Lac-1** and **Lac-2**). However, the much lower yield of this probe made this route less appealing for more precious glycan starting materials, and subsequent conjugations were conducted using **1**.

**Figure 3.**
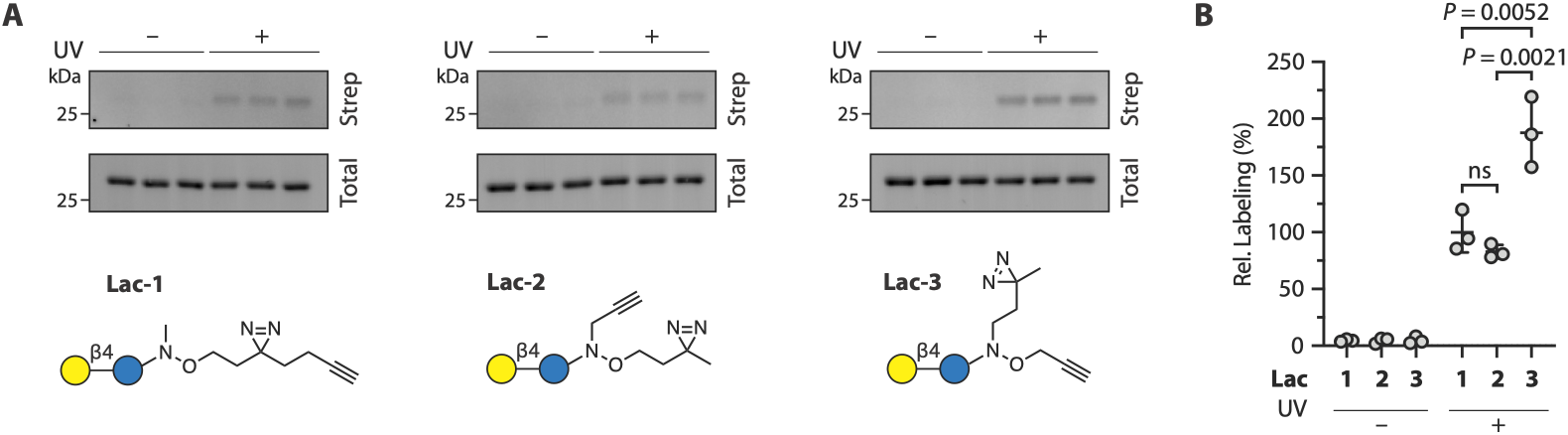
**(A)** Streptavidin blotting and **(B)** quantification of photocrosslinking between 100 µM **Lac-1, Lac-2**, or **Lac-3** probes and LTL. Blots show *n* = 3 independent replicates. Quantification shown as individual replicates with mean ± standard deviation. Samples analyzed using an ordinary one-way ANOVA of +UV samples with Tukey’s multiple comparisons post-hoc test. ns = not significant.

**Scheme 3.**
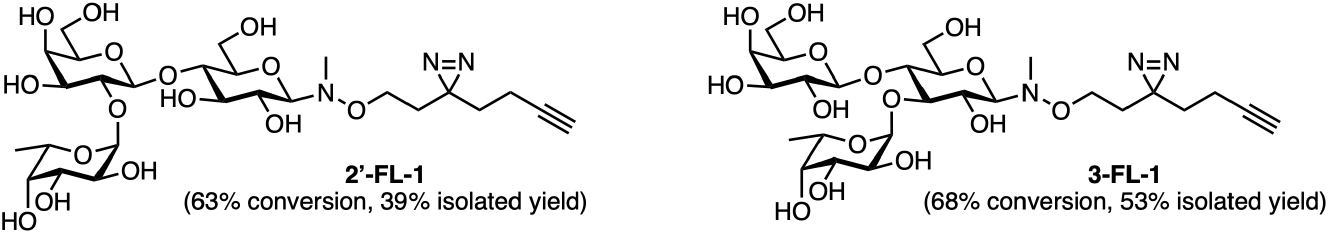
**2’-FL-1** and **3-FL-1** probes.

To show that our minimal probes could maintain selective receptor binding, we synthesized two additional probes based on the human milk oligosaccharides 2’-fucosyllactose (2’-FL) and 3-fucosyllactose (3-FL) using the optimized conjugation condition (**Scheme 3**). ^1^H NMR spectroscopic analysis of reaction crude mixtures revealed that the resulting HMO probes, **2’-FL-1** and **3-FL-1**, were produced in 63% and 68% conversion, respectively, similar to what was observed for **Lac-1**. Purification by organic extraction and SPE columns provided corresponding HMO probes, **2’-FL-1** and **3-FL-1**, in 39% and 53% isolated yield, respectively.

We then subjected these two probes along with **Lac-1** to photocrosslinking experiments with a panel of lectins with known binding affinities. As expected, labeling was strongly dependent on UV irradiation across all samples. Relatively robust labeling was seen for SBA with all three Gal-containing glycans (**Figure 4a**). Although fucosylation has been reported as an impediment for SBA binding,^36^ stronger labeling of SBA was observed for **2’-FL-1** and **3-FL- 1** over **Lac-1**. Labeling was observed from **Lac-1** and **2’-FL-1** to *Erythrina cristagalli lectin* (ECL) (**Figure 4b**), consistent with crystallographic studies supporting its binding activity to lactose and 2’-fucosyllactose sugars.^37, 38^ In addition, labeling of ECL by **3’-FL-1** suggests that its binding cleft also tolerates the incorporation of a 1,3-fucosylation of the lactose scaffold. As expected, photocrosslinking to LTL was seen most strongly for **3-FL-1** (**Figure 4c**), confirming its reported binding affinity to Lewis^X-^like structures. This pattern of selectivity was reversed for *Ulex europaeus* agglutinin I (UEA-I) (**Figure 4d**), which had detectable labeling only for **2’- FL-1**, which contains its preferred recognition motif, Fucα1-2Gal.^36^

**Figure 4.**
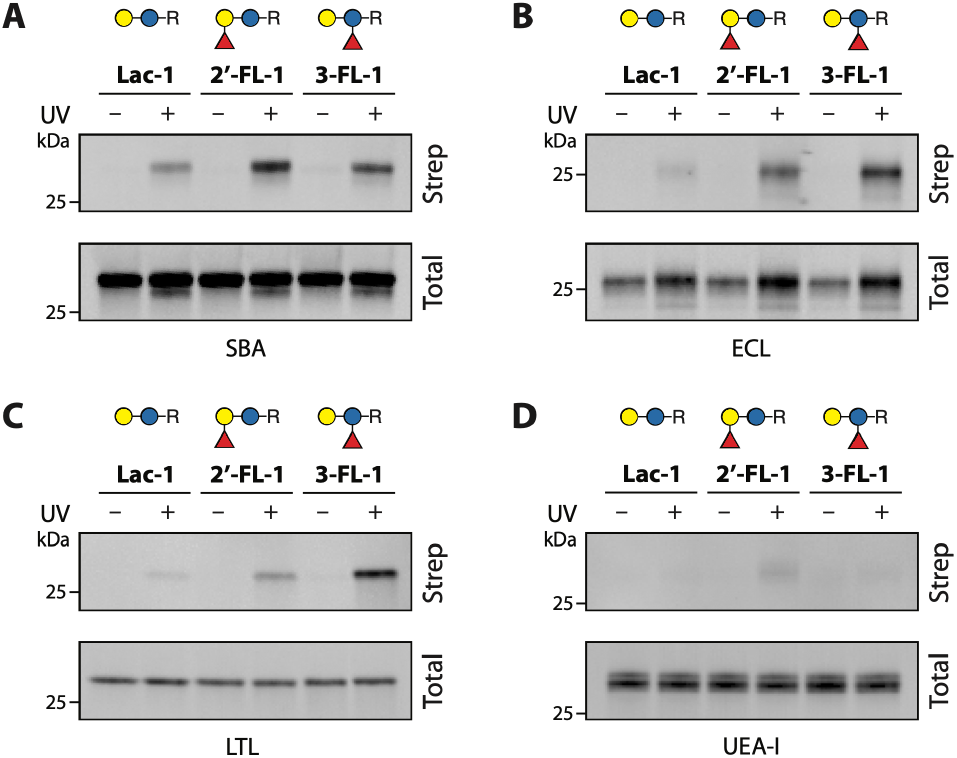
Photocrosslinking between 100 µM **Lac-1, 2’-FL-1**, or **3-FL-1** and lectins **(A)** SBA, **(B)**ECL, **(C)** LTL, and **(D)** UEA-I.

HMOs are known to modulate the intestinal environment in early life through multiple mechanisms, including selective feeding of specific commensal microbiota and inhibition of gut wall adhesion and biofilm formation by opportunistic pathogens.^12, 13^ Moreover, recent evidence has shown that HMOs may also promote healthy aging in adults by altering microbiota composition.^39^ These mechanisms imply that direct interactions between HMOs and microbiota are crucial determinants in their overall beneficial effects. Therefore, we hypothesized that our probes would allow us to directly detect these transient interactions (**Figure 5a**). To show this, we harvested healthy murine fecal material and produced a single cell microbial suspension. We then incubated cells with our probes for 30 min followed by UV irradiation and on-cell CuAAC labeling with AZDye 647 azide. Labeling was then quantified by analytical flow cytometry. Here, we found that both **Lac-1** and **2’-FL-1** strongly labeled microbiota in a concentration-dependent manner (**Figures 5b,c**). However, **3-FL-1** showed minimal labeling even at the highest concentration tested, indicating that direct binding of our glycan probes to microbes within this population was structure-dependent.

**Figure 5.**
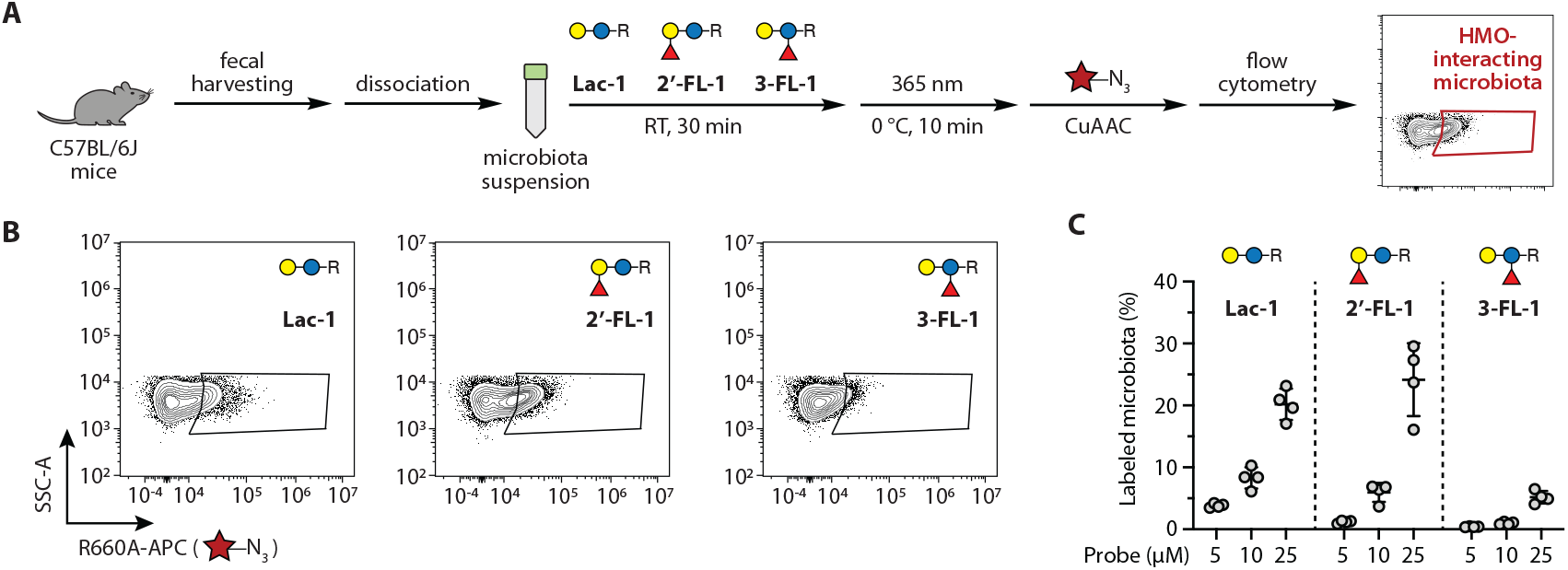
Selective microbiota labeling by **Lac-1, 2’-FL-1**, and **3-FL-1. (A)** Workflow for microbiota labeling and flow cytometric analysis. **(B)** Representative flow cytometry plots and **(C)** quantification of microbiota labeling by photoaffinity HMO probes. *n* = 4 labeling replicates.

Inspired by this high level of photocrosslinking in our population, we aimed to identify these microbes that interacted with **2’-FL-1**. Chemical PRobe INcorporation for Targeted sequencing or PRINT-seq has been employed by several groups to characterize functional microbes within larger communities based on their ability to interact with specific small molecule or biopolymer probes.^40^ For example, metabolic labeling with alkynyl L- or D-amino acids,^41, 42^ noncovalent incorporation of alkynyl cholesterol or fluorescent polysaccharide derivatives,^43, 44^ and enzymatic reactions with alkynyl or fluorescent glycan activity-based probes^45, 46^ have all been utilized to selectively label interacting microbes, which can then be enriched by cell sorting and identified by sequencing. However, this approach has not been utilized with photocrosslinking probes thus far. Therefore, we performed a similar labeling experiment using fecal microbiota from several mice and utilized cell sorting to isolate AZDye 647^+^ microbes for downstream identification (**Figure 6A**).

**Figure 6.**
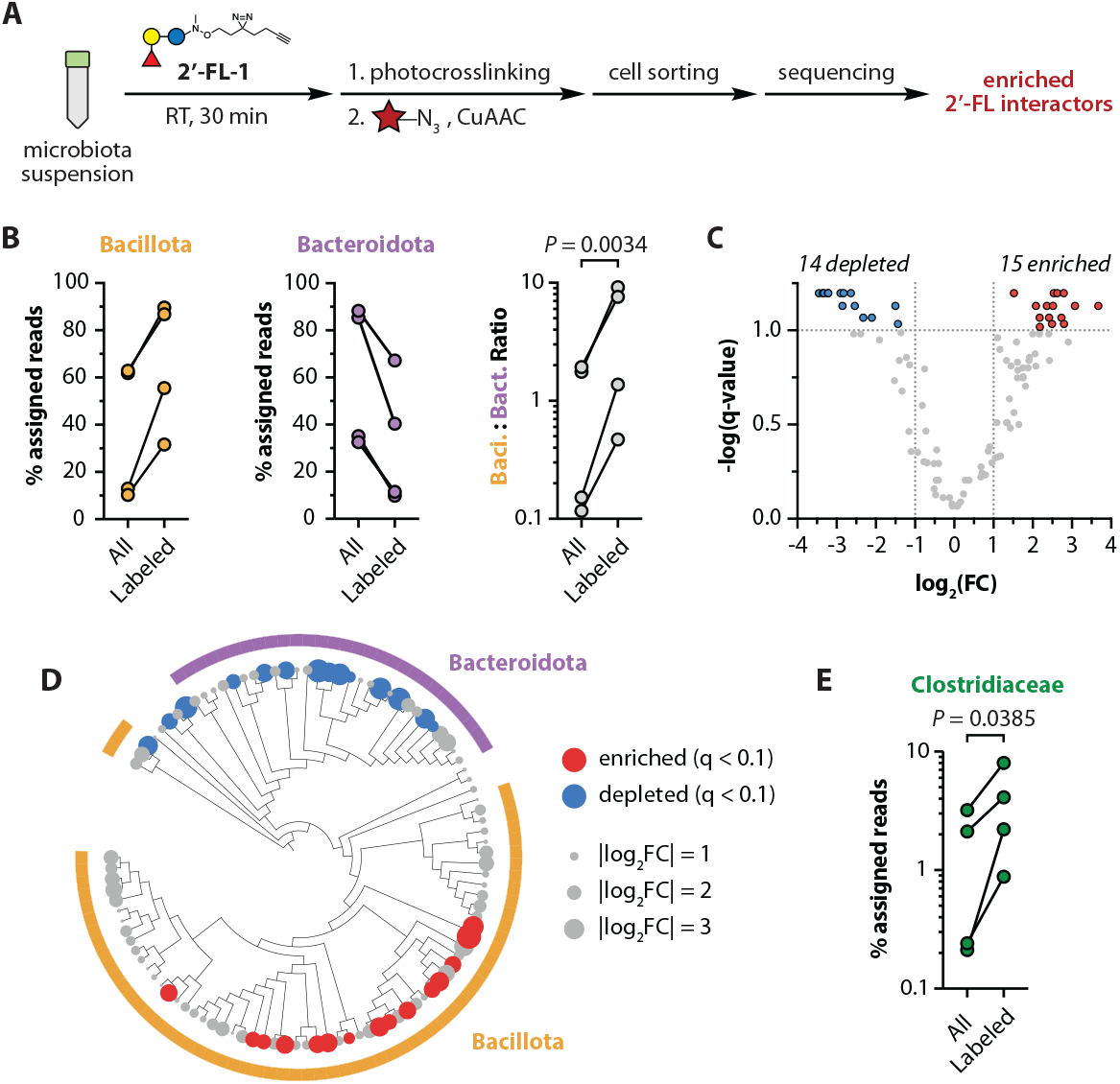
**(A)** Workflow for microbiota labeling, fluorescence-assisted cell sorting, and metagenomic sequencing to identify **2’-FL-1** interactors. **(B)** Changes in percentage of assigned reads to the Bacillota (Firmicutes) and Bacteroidota phyla, and quantification of the Bacillota:Bacteroidota ratio of assigned reads across paired “all” and “labeled” samples. **(C)** Volcano plot and **(D)** cladogram of depleted (blue) and enriched (red) species-level taxa from assigned reads filtered to >0.05% prevalence in four or more samples. **(E)** Changes in percentage of assigned reads for the Clostridiaceae family. Samples in **6B** and **6E** analyzed using a ratio paired *t* test, *n* = 4 mice. Significant species-level taxa in **6C** and **6D** defined as log_2_(fold change) > 1 and *q* < 0.1, as determined by paired *t* tests for each taxon with false discovery rate correction by the Benjamini, Krieger, and Yekutieli method.

Shotgun metagenomic sequencing followed by taxonomic identification revealed significant differences between the total (“all”) and the AZDye 647^+^ (“labeled”) populations (**Tables S4, S5**). Despite high variability between individual mice, all samples demonstrated a significant increase in Bacillota (also known as Firmicutes) and a corresponding decrease in Bacteroidota (**Figure 6B**). When quantified across samples, the ratio between these phyla increased on average by 5.2-fold, which indicated that our approach revealed significant phyla-specific differences in **2’-FL-1** binding. Using normalized read counts, we were able to quantify 105 species-level taxa at or above 0.05% prevalence in four or more samples. We found that 29 taxa exhibited significant depletion or enrichment after targeted sorting (14 depleted and 15 enriched; |fold change| > 2, *q* < 0.1, **Figure 6C**). These changes clustered strongly by phyla (**Figure 6D**), with 15 of 15 enriched taxa found within Bacillota (*P* = 9.8 × 10^- 4^, two-sided Fischer’s exact test) and 11 of 14 depleted taxa found within Bacteroidota (*P* = 2.8 × 10^-5^, two-sided Fischer’s exact test). Recent work has shown that HMOs can be metabolized by commensal clostridia to inhibit the growth of pathobionts.^47^ Corroborating these data, we found an enrichment of the family Clostridiaceae within our AZDye 647^+^ population (**Figure 6E**), indicating that our approach could be useful to identify new beneficial interactions between dietary glycans and gut microbes.

## Conclusion

Herein, we have developed a facile workflow for the one-step synthesis of complex glycan photoaffinity probes. Our newly designed, minimalist linker **1** can be produced using straightforward chemistry, and its conjugation to sugars does not require specialized reagents or equipment. Together, these characteristics provide an appealing and widely accessible strategy to generate complex oligosaccharide probes. Compared to prior generations of linkers using NODHA reactivity, **1** is small and adds less mass to the probe than a single additional hexose unit, minimizing its effects on native glycan binding. Prior work using NODHA compounds have often employed 5-100 equivalents of the linker or excess sugar to drive forward the reaction. Conversely, we were able to achieve NMR conversion values of nearly 70% for more complex oligosaccharides with only four equivalents of **1** using the reducing sugar as the limiting reactant, which greatly simplifies downstream purification.

Importantly, these yields were on par with existing reported methods to generate anomeric glycan photoaffinity probes without the use of excess amounts of the oligosaccharide starting material, making this strategy amenable to more precious glycan starting materials. Synthesis of split linkers **2** and **3** and their corresponding lactose probes highlighted that the coupling efficiency for NODHA compounds is highly influenced by steric hindrance around the nitrogen nucleophile. Nevertheless, the improved crosslinking efficiency seen with **Lac-3** suggests that alternative linker structures in which the diazirine closer to the sugar may further improve target labeling.

Synthesis of and crosslinking experiments with a panel of HMO probes revealed that our linker preserves structure-dependent interactions of glycan-binding proteins, underscoring the minimal nature of our appended linker. Application of our HMO probes to label a complex microbiota community uncovered similar structure-dependent differences in crosslinking, which indicates that our probes can report on selective interactions of live microorganisms with these molecules. The combination of our 2’-FL probe with the PRINT-seq workflow revealed dramatic preferences in **2’-FL-1** binding for Bacillota as well as an enrichment of Clostridiaceae in the labeled fraction of sorted microbiota. Our findings support recent reports of HMO interactions with these commensal microbes and reinforce the idea that HMO interactions with resident microbiota extend beyond the well-characterized interactions with *Bifidobacterium* species. Moreover, our success using these probes to label complex microbiota highlights new opportunities to create glycan probes that will reveal how sugars mediate interactions between the host, diet, and resident microbes.

## Contributions

YC: Investigation, methodology, visualization, writing – original draft, writing – review and editing.

JTL: Methodology, writing – review and editing.

MEG: Conceptualization, formal analysis, investigation, funding acquisition, supervision, visualization, writing – original draft, writing – review and editing.

## Conflicts of interest

The authors declare no competing interests.

## Data availability

Experimental data and procedures are provided in the main text and in the supplementary information. Raw sequencing files are available through BioProject PRJNA1545266. Analytical flow cytometry data are available through Dryad at DOI: 10.5061/dryad.qv9s4mwwv.

## Acknowledgments

This work was supported by the Office of the Director and the National Center for Complementary and Integrative Health (NCCIH) of the National Institutes of Health (NIH) under Award Number DP2AT013749, the National Cancer Institute (NCI) of the NIH under Award Number K22CA272915, the National Science Foundation (NSF GRFP) under Award Number DGE-2235784, and the University of California, Irvine (UCI). The content is solely the responsibility of the authors and does not necessarily represent official views of the NIH or NSF. We thank Samuel Pierce for support in initial fecal collection. We acknowledge the support of the UCI Chemistry Nuclear Magnetic Resonance Facility and the UCI Mass Spectrometry Facility for small molecule characterization. We thank the Prof. Brian Paegel for providing access to the Novocyte flow cytometer.

## Notes

### Competing Interest Statement

The authors have declared no competing interest.

