## Supplemental Information for "One-step glycan photoaffinity probe synthesis enables identification of human milk oligosaccharide-interacting microbiota"

### Supplementary Tables

**Table S1.** Initial attempts to conjugate linker **1** onto lactose.

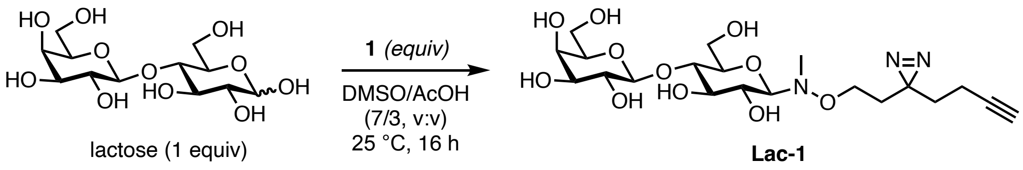

| Entry | **1** (equiv) | Conversion (%, $^1$H NMR)[1] |
|---|---|---|
| 1 | 0.5 | 11 |
| 2 | 1.2 | 22 |
| 3 | 1.5 (4 h) | 38 |
| 4 | 2.0 | 34 |

1. Conversion values were determined by comparing $^1$H NMR integral values for peaks corresponding to the newly formed *N*-glycoside and the remaining reducing sugar in crude reaction mixtures.

**Table S2.** Detailed calculations for stability profiling of diazirine linker in **Table 1**.

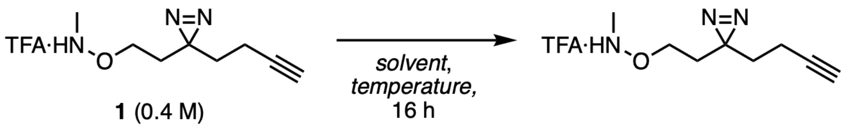

| Entry | Deuterated Solvent | Temp (°C) | Linker:DMS (initial ratio) | Linker:DMS (resulting ratio) | **1** remaining (%)[1] |
|---|---|---|---|---|---|
| 1 | DMSO/AcOH (7:3 v/v) | 25 | 0.435 | – | 0 |
| 2 | | 40 | 0.435 | – | 0 |
| 3 | 1 M NaOAc pH 5 | 25 | 0.435 | 0.417 | 96 |
| 4 | | 40 | 0.435 | 0.345 | 79 |
| 5 | | 65 | 0.392 | – | 0 |
| 6 | Dioxane/NH4OAc (1:1 v/v) pH 5.5 | 25 | 0.435 | 0.395 | 91 |
| 7 | | 40 | 0.435 | 0.345 | 79 |
| 8 | | 65 | 0.392 | 0.108 | 28 |

1. The amount of remaining starting material was calculated as the final ratio divided by the initial ratio of linker to dimethylsulfone (DMS).

**Table S3.** Additional conditions screened for 5-MA–mediated conjugation.

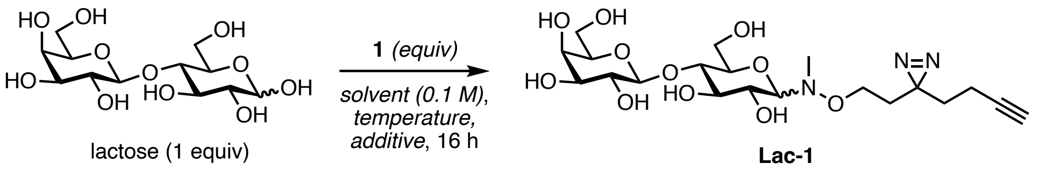

| Entry | Solvent | Temp (°C) | **1** (equiv) | 5-MA (equiv)[1] | Conversion (%)[2] | dr (β:α)[3] |
|---|---|---|---|---|---|---|
| 1 | 1 M NaOAc pH 5.0[4] | 25 | 4.0 | 1.0 | 80 | > 99:1 |
| 3 | | 25 | 4.0 | 1.3 | 79 | > 99:1 |
| 4 | | 40 | 4.0 | 0.7 | 82 | 7:1 |
| 6 | | 40 | 4.0 | 1.0 | 81 | 5.8:1 |

1. 5-MA = 5-methoxyanthranilic acid.
2. Conversion values were determined by comparing $^1$H NMR integral values for peaks corresponding to the newly formed *N*-glycoside and the remaining reducing sugar in crude reaction mixtures.
3. dr = diastereomeric ratio.
4. Buffers were prepared at the indicated pH values and decreased slightly after addition of **1** as the TFA salt.

**Additional tables available as SI Information attachment (.xlsx).**

**Table S4.** Raw assigned read counts.

**Table S5.** Filtered and normalized assigned read counts.

**Table S6.** Graph values.

#### Supplementary Figures

**Figure S1.** Mechanism of NODHA conjugation under acidic conditions.

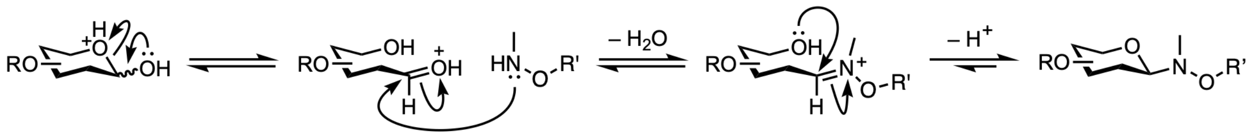

**Figure S2.** Representative data for quantitative microbiota flow cytometric analysis.

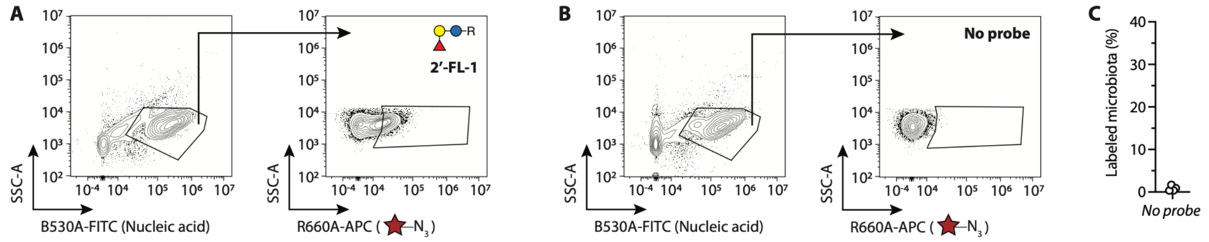

Representative gating strategy for microbiota samples treated with **(A)** 25 µM **2′-FL-1** or **(B)** no probe as described in **Figure 5**. **(C)** Quantification of CuAAC-labeled microbiota treated with no probe.

**Figure S3.** Representative data for microbiota fluorescence-assisted cell sorting.

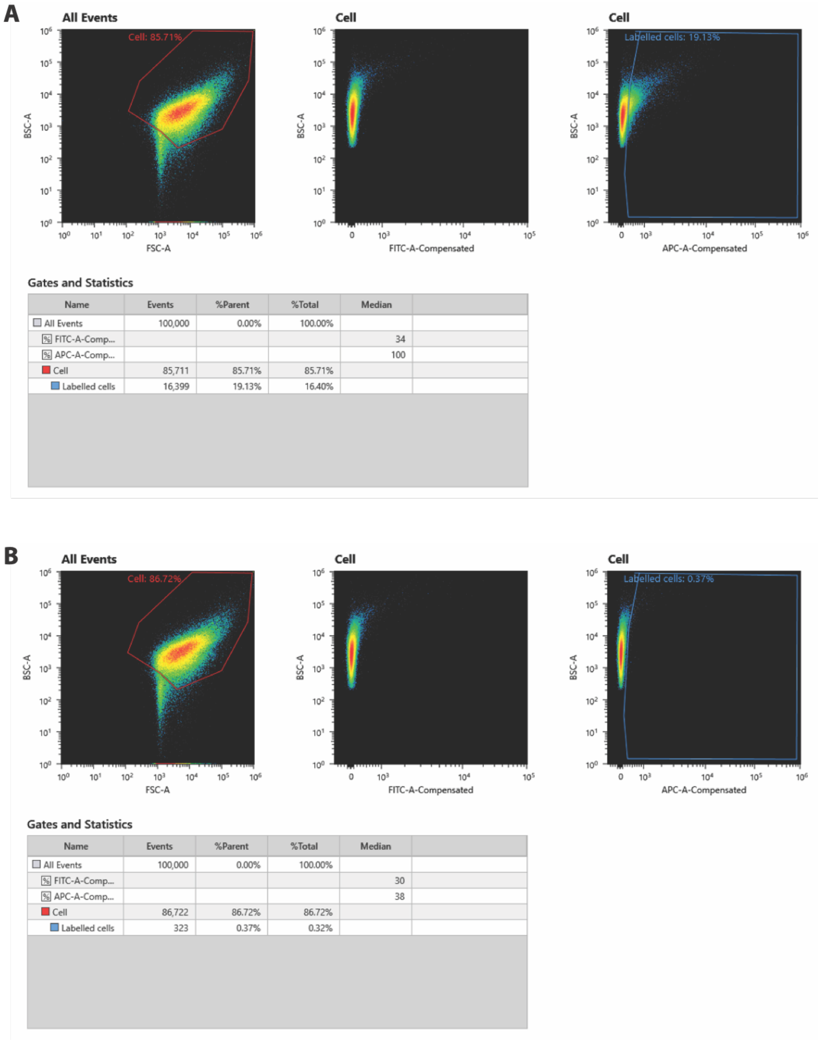

| Name | Events | %Parent | %Total | Median |
|---|---|---|---|---|
| All Events | 100,000 | 0.00% | 100.00% | |
| FITC-A-Comp... | | | | 34 |
| APC-A-Comp... | | | | 100 |
| Cell | 85,711 | 85.71% | 85.71% | |
| Labelled cells | 16,399 | 19.13% | 16.40% | |

| Name | Events | %Parent | %Total | Median |
|---|---|---|---|---|
| All Events | 100,000 | 0.00% | 100.00% | |
| FITC-A-Comp... | | | | 30 |
| APC-A-Comp... | | | | 38 |
| Cell | 86,722 | 86.72% | 86.72% | |
| Labelled cells | 323 | 0.37% | 0.32% | |

Representative gating strategy and quantification for microbiota samples treated with **(A)** 25 µM **2'-FL-1** or **(B)** no probe followed by cell sorting as described in **Figure 6**. Cells were distinguished from debris based on location within FSC-A vs. BSC-A, and labeled cells (AzDye 647$^{+}$) were identified based on intensity in the APC-A channel.

**Figure S4.** Uncropped blots from **Figure 2**.

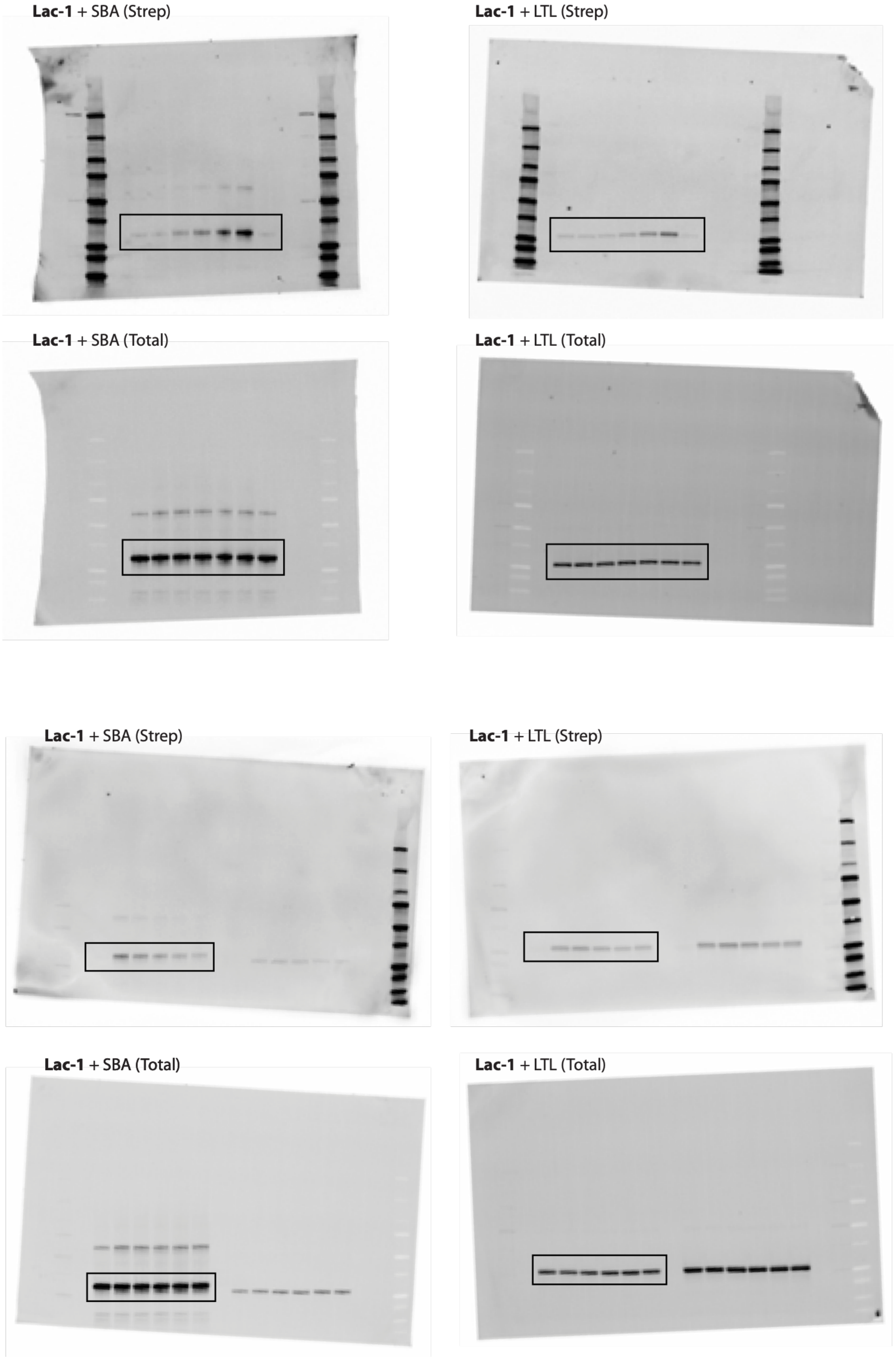

**Figure S5.** Uncropped blots from **Figure 3**.

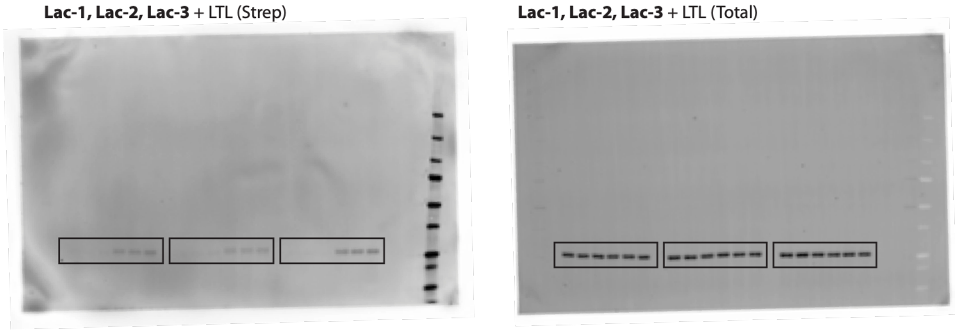

**Figure S6.** Uncropped blots from **Figure 4**.

**Lac-1, 2′-FL-1, 3-FL-1** + SBA (Strep)

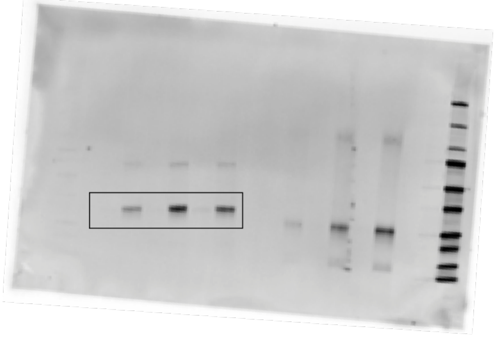

**Lac-1, 2′-FL-1, 3-FL-1** + SBA (Total)

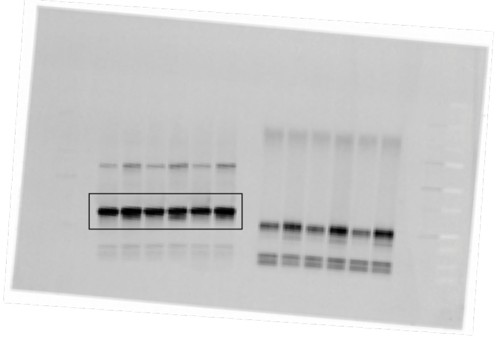

**Lac-1, 2′-FL-1, 3-FL-1** + ECL (Strep)

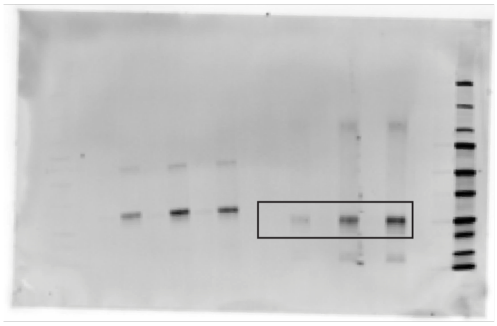

**Lac-1, 2′-FL-1, 3-FL-1** ECL (Total)

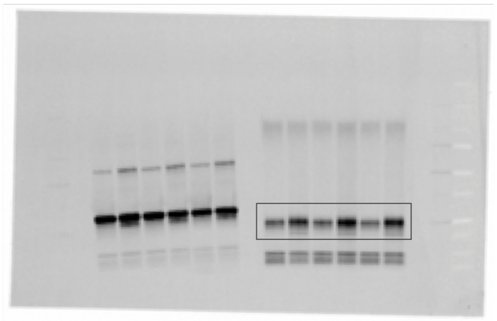

**Lac-1, 2′-FL-1, 3-FL-1** + LTL (Strep)

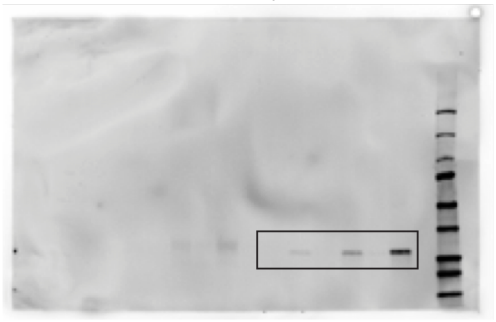

**Lac-1, 2′-FL-1, 3-FL-1** + LTL (Total)

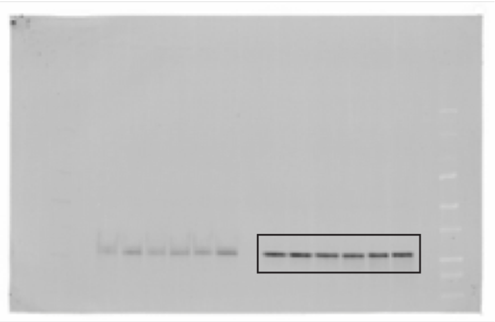

**Lac-1, 2′-FL-1, 3-FL-1** + UEA-I (Strep)

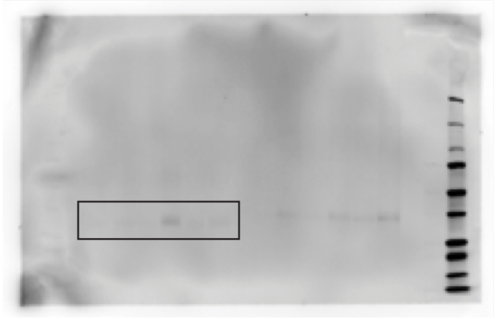

**Lac-1, 2′-FL-1, 3-FL-1** + UEA-I (Total)

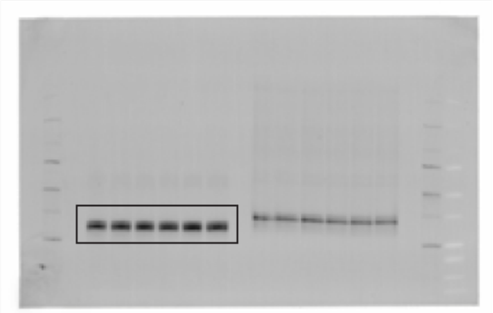

### Materials and Methods

#### *General chemical methods*

Chemical reagents were purchased from Millipore Sigma, Fisher Scientific, VWR, Ambeed, or Chemscene and used as received. Human milk oligosaccharides 2′-FL and 3-FL were obtained from Elicityl at 95% purity and used as provided. All chemical reactions were performed in flame-dried or oven-dried glassware under $N_2$. Organic solvents including dichloromethane and acetonitrile were commercial dry solvents or freshly dried solvents passed through an activated alumina column under Ar. All solvents used were ACS reagent or HPLC grade. All aqueous buffers were prepared using Milli-Q $H_2O$. Reactions were monitored by thin layer chromatography (TLC) using SiliCycle SiliaPlate glass-backed plates (250 µm particle size; 01-184-505, Fisher Scientific). Automated flash chromatography was accomplished using a Teledyne Isco CombiFlash NextGen 300+ system using RediSep Gold silica gel flash columns (20-40 µm particle size, Teledyne Isco). $^{1}H$ and $^{13}C\{^{1}H\}$ nuclear magnetic resonance (NMR) experiments were recorded on Bruker Avance spectrometers at 500 MHz for $^{1}H$ spectra and 126 MHz for $^{13}C$ spectra at the UC Irvine Chemistry NMR Facility. Data for $^{1}H$ NMR spectra are reported as follows: chemical shift relative to the deuterated solvent (δ ppm), multiplicity, coupling constant (Hz), and integration. Data for $^{13}C$ NMR spectra are reported as chemical shift. All NMR data are reported based on calibration to the chemical shifts of the following deuterated solvents: $^{1}H$ NMR: $CDCl_3$ $\delta$ 7.26 ppm; $D_2O$ $\delta$ 4.76 ppm; $CD_3OD$ $\delta$ 3.31 ppm. $^{13}C$ NMR: $CDCl_3$ $\delta$ 77.16 ppm; $CD_3OD$ $\delta$ 49.00 ppm. Multiplicity is reported as follows: s = singlet, br s = broad singlet, d = doublet, ddt = doublet of doublet of triplet, t = triplet, dd = doublet of doublet, dt = doublet of triplet, m = multiplet. High-resolution mass spectra (HRMS) were recorded on a Waters Micromass LCT Premier at the UC Irvine Chemistry Mass Spectrometry Facility.

#### *Synthesis of N-Me-O-DiazAlk 1*

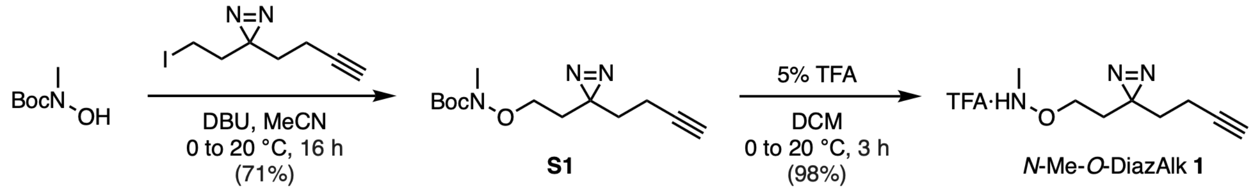

**_tert_-Butyl (2-(3-(but-3-yn-1-yl)-3*H*-diazirin-3-yl)ethoxy)(methyl)carbamate (S1).**

In MeCN (10 mL), 3-(but-3-yn-1-yl)-3-(2-iodoethyl)-3H-diazirine (0.595 g, 4.04 mmol, 2.1 equiv) and *tert*-butyl hydroxy(methyl)carbamate (0.470 g, 1.89 mmol, 1.0 equiv) was dissolved and cooled to 0 °C. DBU (0.603 mL, 4.04 mmol, 2.1 equiv) was then added dropwise, and the reaction was stirred at 0 °C for an additional 20 min. The reaction was allowed to warm to 20 °C and continue to stir overnight. After 16 h, the reaction mixture was diluted with Milli-Q $H_2O$ (10 mL). The mixture was extracted with $CH_2Cl_2$ (2 × 10 mL). The combined organic layers were washed with brine (20 mL), dried over $Na_2SO_4$, filtered, and concentrated *in vacuo*. Purification by flash column chromatography (19:1 hexanes/EtOAc) afforded carbamate **S1** (0.361 g, 71%) as a yellow oil.

$^{1}$H NMR (500 MHz, $CDCl_3$) $\delta$ 3.69 (t, $J$ = 6.4 Hz, 2H), 3.09 (s, 3H), 2.03 (td, $J$ = 7.5, 2.6 Hz, 2H), 1.98 (t, $J$ = 2.7 Hz, 1H), 1.73 (t, $J$ = 6.3 Hz, 2H), 1.69 (t, $J$ = 7.5 Hz, 2H), 1.49 (s, 9H).

$^{13}$C NMR (126 MHz, $CDCl_3$) $\delta$ 157.0 (C), 82.9 (C), 81.6 (C), 69.3 (CH), 68.6 ($CH_2$), 36.9 ($CH_3$), 32.7 ($CH_2$), 31.8 ($CH_2$), 28.4 ($CH_3$), 27.8 (C), 13.4 ($CH_2$).

HRMS (TOF MS ES$^+$) *m/z* calculated for $C_{13}H_{21}N_3O_3Na$ $(M + Na)^+$ 290.1481, found 290.1490.

**_O_-(2-(3-(But-3-yn-1-yl)-3_H_-diazirin-3-yl)ethyl)-_N_-methylhydroxylamine TFA salt (_N_-Me-_O_-DiazAlk 1).**

Carbamate **S1** (0.360 g, 1.35 mmol) was dissolved in $CH_2Cl_2$ (34 mL) and cooled to 0 °C. TFA (20% in $CH_2Cl_2$, 11 mL) was added dropwise, and the reaction was stirred at 0 °C for an additional 20 min. The reaction was then allowed to warm to 20 °C. After 3 h, the reaction mixture was concentrated in *vacuo* to afford the TFA salt of *N*-Me-*O*-DiazAlk **1** (0.388 g, 98%) as a yellow oil, which was used without further purification.

$^{1}$H NMR (500 MHz, DMSO-$d_6$) $\delta$ 3.79 (t, $J$ = 6.4 Hz, 2H), 2.85 (t, $J$ = 2.6 Hz, 1H), 2.82 (s, 3H), 2.03 (td, $J$ = 7.4, 2.7 Hz, 2H), 1.75 (t, $J$ = 6.4 Hz, 2H), 1.63 (t, $J$ = 7.5 Hz, 2H).

$^{13}$C NMR (126 MHz, DMSO-$d_6$) $\delta$ 83.6 (C), 72.3 (CH), 68.0 ($CH_2$), 36.1 ($CH_3$), 31.9 ($CH_2$), 31.2 ($CH_2$), 27.1 (C), 13.4 ($CH_2$). HRMS (TOF MS ES$^+$) *m/z* calculated for $C_8H_{13}N_3O$ $(M)^+$ 167.1059, found 167.1080.

A fragment arising from N–O bond cleavage was also detected:
HRMS (TOF MS ES$^+$) *m/z* calculated for $C_7H_9N_2OH$ $(M + H)^+$ 138.0793, found 138.0799.

***Synthesis of Boc-protected N-Alk-O-Diaz (Boc-2)***

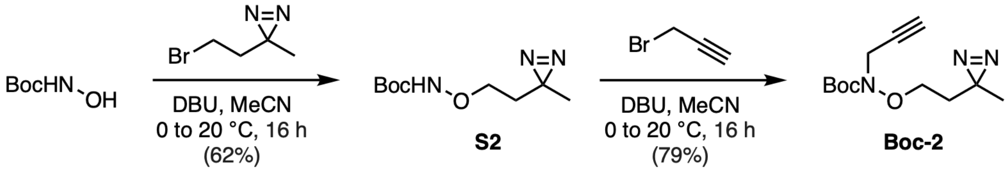

**_tert_-Butyl (2-(3-methyl-3_H_-diazirin-3-yl)ethoxy)carbamate (S2).**

In MeCN (3 mL), 3-(2-bromoethyl)-3-methyl-3*H*-diazirine (0.100 g, 0.613 mmol, 1.0 equiv) and *tert*-butyl hydroxycarbamate (0.106 g, 0.798 mmol, 1.3 equiv) was dissolved and cooled to 0 °C. DBU (0.119 mL, 0.798 mmol, 1.3 equiv) was then added dropwise, and the reaction was stirred at 0 °C for an additional 20 min. The reaction was allowed to warm to 20 °C and continue to stir overnight. After 16 h, the reaction mixture was diluted with Milli-Q $H_2O$ (5 mL). The mixture was extracted with $CH_2Cl_2$ (2 × 5 mL). The combined organic layers were washed with brine (10 mL), dried over $Na_2SO_4$, filtered, and concentrated *in vacuo*. Purification by flash column chromatography (9:1 hexanes/EtOAc) afforded carbamate **S2** (0.0824 g, 62%) as a yellow oil.

$^{1}$H NMR (500 MHz, $CDCl_3$) δ 7.17 (br s, 1H), 3.80 (t, $J$ = 6.9 Hz, 2H), 1.64 (t, $J$ = 6.2 Hz, 2H), 1.48 (s, 9H), 1.06 (s, 3H).

$^{13}C$ NMR (126 MHz, $CDCl_3$) $\delta$ 157.1 (C), 82.1 (C), 71.5 ($CH_2$), 33.3 ($CH_2$), 28.3 ($CH_3$), 24.2 (C), 20.0 ($CH_3$).

HRMS (TOF MS ES$^+$) *m/z* calculated for $C_9H_{17}N_3O_3Na$ $(M + Na)^+$ 238.1168, found 238.1167.

**_tert_-Butyl (2-(3-methyl-3*H*-diazirin-3-yl)ethoxy)(prop-2-yn-1-yl)carbamate (Boc-2).**

In MeCN (2 mL), propargyl bromide (0.0535 g, 0.440 mmol, 1.3 equiv) and carbamate **S2** (0.0731 g, 0.340 mmol, 1.0 equiv) was dissolved and cooled to 0 °C. DBU (0.0660 mL, 0.440 mmol, 1.3 equiv) was then added dropwise, and the reaction was stirred at 0 °C for an additional 20 min. The reaction was allowed to warm to 20 °C and continue to stir overnight. After 16 h, the reaction mixture was diluted with $CH_2Cl_2$ (3 mL) and Milli-Q $H_2O$ (5 mL). The mixture was extracted with $CH_2Cl_2$ (2 × 5 mL). The combined organic layers were washed with brine (10 mL), dried over $Na_2SO_4$, filtered, and concentrated *in vacuo*. Purification by flash column chromatography (9:1 hexanes/EtOAc) afforded carbamate **Boc-2** (0.0682 g, 79%) as a yellow oil.

$^1H$ NMR (500 MHz, $CDCl_3$) $\delta$ 4.19 (d, $J$ = 2.6 Hz, 2H), 3.87 (t, $J$ = 6.5 Hz, 2H), 2.24 (t, $J$ = 2.3 Hz, 1H), 1.66 (t, $J$ = 6.3 Hz, 2H), 1.50 (s, 9H), 1.08 (s, 3H).

$^{13}C$ NMR (126 MHz, $CDCl_3$) $\delta$ 156.5 (C), 82.6 (C), 78.5 (C), 71.9 ($CH_2$), 70.5 (CH), 40.4 ($CH_2$), 33.5 ($CH_2$), 28.3 ($CH_3$), 24.3 (C), 20.3 ($CH_3$).

HRMS (TOF MS ES$^+$) *m/z* calculated for $C_{12}H_{19}N_3O_3Na$ $(M + Na)^+$ 276.1324, found 276.1328.

***Synthesis of Boc-protected N-Diaz-O-Alk (Boc-3)***

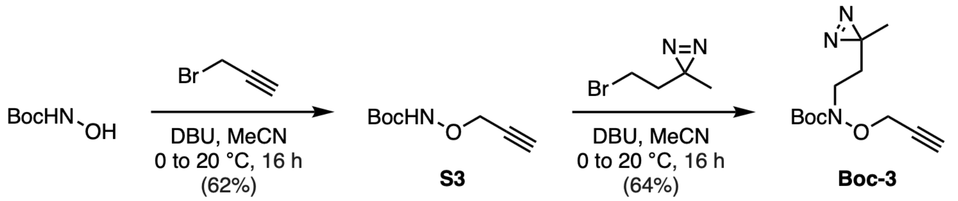

**_tert_-Butyl (prop-2-yn-1-yloxy)carbamate (S3).**

In MeCN (25 mL), propargyl bromide (1.00 g, 8.41 mmol, 1.0 equiv) and *tert*-butyl hydroxycarbamate (1.46 g, 10.9 mmol, 1.3 equiv) was dissolved and cooled to 0 °C. DBU (1.63 mL, 10.9 mmol, 1.3 equiv) was then added dropwise, and the reaction was stirred at 0 °C for an additional 20 min. The reaction was allowed to warm to 20 °C and continue to stir overnight. After 16 h, the reaction mixture was diluted with Milli-Q $H_2O$ (25 mL). The mixture was extracted with $CH_2Cl_2$ (2 × 25 mL). The combined organic layers were washed with brine (25 mL), dried over $Na_2SO_4$, filtered, and concentrated *in vacuo*. Purification by flash column chromatography (9:1 hexanes/EtOAc) afforded carbamate **S3** (2.24 g, 78%) as a yellow oil.

$^1H$ NMR (500 MHz, $CDCl_3$) $\delta$ 7.33 (br s, 1H), 4.47 (d, $J$ = 2.4 Hz, 2H), 2.50 (t, $J$ = 2.4 Hz, 1H), 1.48 (s, 9H).

$^{13}C$ NMR (126 MHz, $CDCl_3$) $\delta$ 156.5 (C), 82.3 (C), 78.4 (C), 75.7 (CH), 63.9 ($CH_2$), 28.3 ($CH_3$).

HRMS (TOF MS ES+) *m/z* calculated for $C_8H_{13}NO_3Na$ $(M + Na)^+$ 194.0793, found 194.0787.

**_tert_-Butyl (2-(3-methyl-3*H*-diazirin-3-yl)ethyl)(prop-2-yn-1-yloxy)carbamate (Boc-3).**

In MeCN (20 mL), 3-(2-bromoethyl)-3-methyl-3*H*-diazirine (0.250 g, 1.50 mmol) and carbamate **S4** (1.26 g, 7.36 mmol) was dissolved and cooled to 0 °C. DBU (1.10 mL, 7.36 mmol) was then added dropwise, and the reaction was stirred at 0 °C for an additional 20 min. The reaction was allowed to warm to 20 °C and continue to stir overnight. After 16 h, the reaction mixture was diluted with Milli-Q $H_2O$ (20 mL). The mixture was extracted with $CH_2Cl_2$ (2 × 20 mL). The combined organic layers were washed with brine (20 mL), dried over $Na_2SO_4$, filtered, and concentrated *in vacuo*. Purification by flash column chromatography (9:1 hexanes/EtOAc) afforded carbamate **Boc-3** (0.241 g, 64%) as a yellow oil.

$^1$H NMR (500 MHz, $CDCl_3$) δ 4.49 (d, *J* = 2.4 Hz, 2H), 3.52–3.50 (m, 2H), 2.48 (t, *J* = 2.4 Hz, 1H), 1.63–1.61 (m, 2H), 1.50 (s, 9H), 1.04 (s, 3H).

$^{13}$C NMR (126 MHz, $CDCl_3$) *δ* 156.5 (C), 82.2 (C), 78.7 (C), 75.7 (CH), 62.6 ($CH_2$), 45.8 ($CH_2$), 32.1 ($CH_2$), 28.4 ($CH_3$), 24.2 (C), 19.7 ($CH_3$).

HRMS (TOF MS ES+) *m/z* calculated for $C_{12}H_{19}N_3O_3Na$ $(M + Na)^+$ 276.1324, found 276.1325.

##### *Degradation studies of N-Alk-O-Diaz 1*

A solution of *N*-Alk-*O*-Diaz **1** was prepared at 0.4 M in the corresponding deuterated solvent system. Dimethylsulfone (DMS) was added as an internal standard, and the initial ratio of **1** to DMS was determined by $^1$H NMR spectroscopy prior to reaction. The reaction mixture was then stirred at 20, 40, or 65 °C for 16 h and analyzed directly by $^1$H NMR to obtain the final ratio of **1** to DMS. Degradation of **1** was calculated by dividing the final **1**/DMS ratio by the initial **1**/DMS ratio and multiplying by 100.

##### *General procedure for screening conjugation reactions with N-Alk-O-Diaz 1*

To a solution of lactose (1.0 equiv) in the corresponding deuterated buffer (0.1 M) was added the *N*-Alk-*O*-Diaz **1** at the desired equivalents without or with 5-methoxyanthranilic acid at the denoted equivalents. The reaction mixture was then stirred at the desired temperature for 16 h. The reaction crude mixtures were analyzed directly by $^1$H NMR spectroscopy without any further purification. Crude conversions of conjugation reactions were quantified using the Glc H-1 anomeric signals and calculated as the integration of the Glc H-1 signal of **Lac-1** divided by the sum of the integrations of the corresponding Glc H-1 signals of **Lac-1** and unreacted lactose.

##### *General procedure for conjugation reactions of unprotected sugars*

To a solution of unprotected sugar (1.0 equiv, 0.1 M) in 1 M NaOAc pH 5 at room temperature was added the NODHA linker (4.0 equiv). The final pH of the reaction mixture was measured at pH 4.5. The reaction mixture was stirred at room temperature for 16 h. The reaction mixture was then diluted with Milli-Q $H_2O$ (10 mL per mmol of unprotected sugar) and extracted with $CH_2Cl_2$ (20 mL per mmol of unprotected sugar). The separated organic layer was concentrated *in vacuo* to recover the unreacted NODHA linker as the acetate salt. The aqueous layer was concentrated *in vacuo,* lyophilized to remove residual $H_2O$, and analyzed by $^1$H NMR spectroscopy.

The crude mixture was then redissolved in 1 mL of Milli-Q $H_2O$ for purification by solid phase extraction. For each reaction, an Oasis HLB column (Waters, 186000115; 6 cc vacuum cartridge, 500 mg sorbent per cartridge, 60 µm) was attached to a vacuum manifold system (Waters, WAT200607) and activated with 2 mL of methanol followed by 2 mL of Milli-Q $H_2O$. The crude mixture was then added to the column and passed through at a flow rate corresponding to one drop per second. The column was washed with 2 mL of Milli-Q $H_2O$ twice to remove unreacted sugar. The conjugated product was then eluted using a stepwise gradient of 2-mL elution fractions of 0–40% aqueous acetonitrile. Fractions containing the product were combined, concentrated *in vacuo*, and lyophilized to remove residual $H_2O$.

For reactions using **2** and **3**, the linkers were generated by TFA deprotection and used directly for glycan conjugation. Here, **Boc-2** or **Boc-3** was dissolved in $CH_2Cl_2$ at 40 mM and cooled to 0 °C. A solution of 20% TFA in $CH_2Cl_2$ was added dropwise until the reaction reached a final concentration of 5% TFA. The reaction was then stirred at 0 °C for 20 min and then allowed to warm to 20 °C. After 3 h, the reaction mixture was concentrated *in vacuo* to afford the TFA salt of the respective NODHA linker. Successful deprotection was verified by TLC and HRMS before directly proceeding to the respective conjugation reaction as described above.

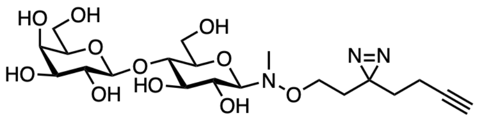

**Lac-1**

**Lac-1.** Scale: 15.0 mg, 0.0483 mmol; Conversion: 61%; Isolated yield: 8.8 mg, 41%.

$^1$H NMR (500 MHz, $CD_3OD$) $\delta$ 4.37 (d, $J$ = 7.6 Hz, 1H), 4.03 (d, $J$ = 8.8 Hz, 1H), 3.90 (td, $J$ = 12.6, 2.5 Hz, 1H), 3.87–3.76 (m, 3H), 3.70 (dd, $J$ = 11.4, 4.7 Hz, 1H), 3.65–3.50 (m, 6H), 3.50–3.43 (m, 2H), 3.40–3.35 (m, 1H), 2.73 (s, 3H), 2.27 (t, $J$ = 3.0 Hz, 1H), 2.03 (td, $J$ = 7.3, 2.7 Hz, 2H ),1.71 (t, $J$ = 5.6 Hz, 2H), 1.66–1.58 (m, 2H).

$^{13}$C NMR (126 MHz, $CD_3OD$) $\delta$ 105.1 (CH), 95.2 (CH), 83.7 (C), 80.5 (CH), 78.2 (CH), 77.7 (CH), 77.1 (CH), 74.8 (CH), 72.6 (CH), 71.4 (CH), 70.3 (CH), 68.3 ($CH_2$), 62.5 ($CH_2$), 62.1 ($CH_2$), 39.1 ($CH_3$), 33.8 ($CH_2$), 32.7 ($CH_2$), 27.8 (C), 13.8 ($CH_2$).

HRMS (TOF MS ES$^+$) $m/z$ calculated for $C_{20}H_{34}N_3O_{11}Na$ $(M + H)^+$ 492.2193, found 492.2193.

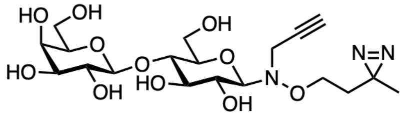

**Lac-2**

**Lac-2.** Scale: 15.0 mg, 0.0483 mmol; Conversion: 19%; Isolated yield: 3.0 mg, 14%.

$^{1}$H NMR (500 MHz, $CD_3OD$) $\delta$ 4.27 (d, $J$ = 7.4 Hz, 1H), 4.09 (d, $J$ = 8.4 Hz, 1H), 3.82–3.66 (m, 7H), 3.62 (td, $J$ = 10.9, 2.8 Hz, 1H), 3.51–3.42 (m, 5H), 3.39 (dd, $J$ = 9.5, 3.3 Hz, 1H), 3.28–3.24 (m, 1H), 2.52 (t, $J$ = 2.6 Hz, 1H), 1.50 (t, $J$ = 6.6 Hz, 1H), 1.22–1.18 (m, 2H), 0.94 (s, 3H).

$^{13}$C NMR (126 MHz, $CD_3OD$) $\delta$ 103.8 (CH), 95.2 (CH), 80.5 (CH), 78.2 (CH), 77.7 (CH), 77.1 (CH), 74.8 (CH), 72.6 (CH), 71.4 (CH), 70.3 (CH), 68.3 ($CH_2$), 62.5 ($CH_2$), 62.1 ($CH_2$), 39.1 ($CH_3$), 33.8 ($CH_2$), 32.7 ($CH_2$), 27.8 (C), 13.8 ($CH_2$).

HRMS (TOF MS ES$^+$) *m/z* calculated for $C_{19}H_{32}N_3O_{11}$ $(M + H)^+$ 478.2037, found 478.2038.

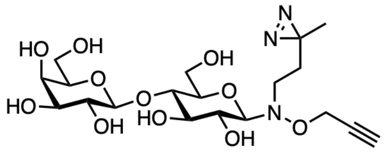

Lac-3

**Lac-3.** Scale: 15.0 mg, 0.0483 mmol; Conversion: 15%; Isolated yield: 2.7 mg, 13%.

$^{1}$H NMR (500 MHz, $CD_3OD$) $\delta$ 4.34 (dd, $J$ = 8.8, 2.4 Hz, 2H), 4.27 (d, $J$ = 7.8 Hz, 1H), 4.00 (d, $J$ = 7.8 Hz, 1H), 3.77 (td, $J$ = 12.4, 2.6 Hz, 1H), 3.60 (dd, $J$ = 11.6, 4.4 Hz, 1H), 3.51–3.36 (m, 7H), 3.27 (dt, $J$ = 9.0, 3.1 Hz, 1H), 2.93–2.87 (m, 1H), 2.83–2.76 (m, 1H), 2.81 (t, $J$ = 2.2 Hz, 1H), 1.24–1.20 (m, 2H), 1.57–1.51 (m, 2H), 0.94 (s, 3H).

$^{13}$C NMR (126 MHz, $CD_3OD$) $\delta$ 103.7 (CH), 92.8 (CH), 78.7 (CH), 76.2 (CH), 75.7 (CH), 75.1 (C), 73.4 (CH), 71.2 (CH), 69.9 (CH), 68.9 ($CH_2$), 61.9 ($CH_2$), 61.2 ($CH_2$), 60.6 ($CH_2$), 32.1 ($CH_2$), 29.2 ($CH_2$), 24.3 (C), 18.5 ($CH_3$).

HRMS (TOF MS ES$^+$) *m/z* calculated for $C_{19}H_{31}N_3O_{11}Na$ $(M + Na)^+$ 500.1324, found 500.1858.

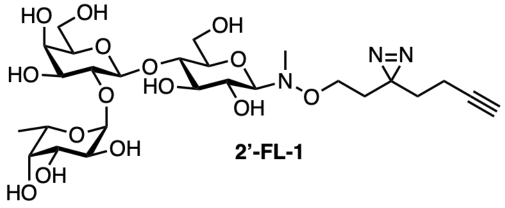

2'-FL-1

**2'-FL-1.** Scale: 13.0 mg, 0.0253 mmol; Conversion: 45%; Isolated yield: 6.3 mg, 39%.

$^{1}$H NMR (500 MHz, $CD_3OD$) $\delta$ 5.28 (d, $J$ = 3.2 Hz, 1H), 4.51 (d, $J$ = 7.5 Hz, 1H), 4.21 (dt, $J$ = 13.2, 6.3 Hz, 1H), 4.09–4.05 (m, 1H), 4.00–3.94 (m, 1H), 3.89–3.82 (m, 2H), 3.82–3.54 (m, 8H), 3.42–3.37 (m, 1H), 2.70 (s, 3H), 2.36 (t, $J$ = 2.4 Hz, 1H), 2.01 (td, $J$ = 7.0, 2.6 Hz, 2H),1.81–1.74 (m, 2H), 1.73–1.66 (m, 2H), 1.20 (d, $J$ = 6.6 Hz, 3H).

$^{13}$C NMR (126 MHz, $CD_3OD$) $\delta$ 102.6 (CH), 101.8 (CH), 98.2 (CH), 83.7 (C), 78.9 (CH), 78.8 (CH), 78.6 (CH), 78.2 (CH), 78.0 (CH), 77.7 (CH), 77.0 (CH), 76.4 (CH), 76.0 (CH), 75.4 (CH), 75.3 (CH), 73.6 (CH), 71.7 (CH), 70.7 ($CH_2$), 68.3 ($CH_2$), 62.5 ($CH_2$), 39.0 ($CH_2$), 33.8 ($CH_2$), 32.7 ($CH_2$), 27.7 (C), 16.7 ($CH_2$), 13.9 ($CH_3$).

HRMS (TOF MS ES$^+$) *m/z* calculated for $C_{26}H_{43}N_3O_{16}Na$ (M + Na)$^+$ 660.2592, found 660.2581.

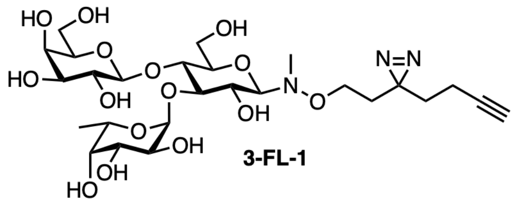

3-FL-1

**3-FL-1.** Scale: 13.0 mg, 0.0253 mmol; Conversion: 68%; Isolated yield: 8.5 mg, 53%.

$^1$H NMR (500 MHz, $CD_3OD$) $\delta$ 5.43 (d, *J* = 4.3 Hz, 1H), 4.43 (d, *J* = 8.0 Hz, 1H), 4.03 (d, *J* = 8.9 Hz, 1H), 3.97–3.90 (m, 3H), 3.86–3.65 (m, 9H), 3.63–3.55 (m, 2H), 3.53–3.43 (m, 3H), 3.36 (dt, *J* = 9.7, 3.1 Hz, 1H), 2.72 (s, 3H), 2.28 (t, *J* = 2.8 Hz, 1H), 2.03 (td, *J* = 7.8, 2.8 Hz, 2H),1.71 (td, *J* = 6.6, 2.1 Hz, 2H), 1.67–1.60 (m, 2H), 1.18 (d, *J* = 6.8 Hz, 3H).

$^{13}$C NMR (126 MHz, $CD_3OD$) $\delta$ 103.8 (CH), 100.4 (CH), 95.3 (CH), 83.7 (C), 80.9 (CH), 79.1 (CH), 76.7 (CH), 74.9 (CH), 74.4 (CH), 73.8 (CH), 73.1 (CH), 72.9 (CH), 71.3 (CH), 70.5 (CH), 70.4 (CH), 70.0 (CH), 68.3 ($CH_2$), 67.5 (CH), 62.9 ($CH_2$), 61.6 ($CH_2$), 39.1 ($CH_3$), 33.8 ($CH_2$), 32.7 ($CH_2$), 27.7 (C), 16.7 ($CH_3$), 13.8 ($CH_2$).

HRMS (TOF MS ES$^+$) *m/z* calculated for $C_{26}H_{43}N_3O_{16}Na$ (M + Na)$^+$ 660.2592, found 660.2610.

##### ***Lectin photocrosslinking***

For lectin labeling experiments, the following lectins were purchased from Vector Labs and used as received: *Lotus tetragonolobus* lectin (LTL, L-1320-5); soybean agglutinin (SBA, L-1010-10); *Ulex europaeus* agglutinin I (UEA-I, L-1060); and *Erythrina cristagalli* lectin (ECL, FL-1141-5). Photocrosslinking experiments were performed at 20 µL scale in PCR tubes using the corresponding glycan probes at indicated concentrations with or without the unmodified glycan along with individual lectins (final concentration: 1 mg mL$^{-1}$). Probes and lectins were prepared in ice-cold PBS immediately prior to photocrosslinking experiments. Samples were gently mixed and then incubated for 20 min at 0 °C in the dark. Samples were then irradiated with 365 nm light for 10 min on ice using a Spectrolinker XL-1000 UV crosslinker (Spectro-UV) with the tubes held at 2 cm from the light source. No-UV negative controls were incubated for 10 min on ice in the dark. Samples were then diluted with 26.5 µL of ice-cold PBS.

CuAAC mix was prepared separately by combining in order 2 parts of 50 mM CuSO4•5$H_2O$, 2 parts of 50 mM TCEP•HCl, 1 part of 10 mM BTTAA, and 1 part of 10 mM biotin-PEG$_3$-azide with mixing between each addition. To each sample, 3.5 µL of the CuAAC mix was added, and the reactions were incubated for 1 h at room temperature in the dark. Samples were then diluted with 10 µL of PBS and 20 µL of 4x Laemmli sample buffer (Bio-Rad, 1610747) and boiled for 10 min. Samples were loaded onto a 4–15% Criterion TGX Midi protein gel (5671084, Bio-Rad) and resolved at 200 V using 1x Tris/SDS/glycine buffer (1610732, Bio-Rad). Total protein loading was measured using Stain-Free imaging on a Bio-Rad ChemiDoc MP Imaging System using the manufacturer's settings. Lectins were then transferred by first assembling a transfer sandwich using a 0.45 µm low-fluorescence PVDF membrane from the Trans-Blot Turbo RTA Midi transfer kit (1704275, Bio-Rad). Transfer assemblies were then run on a Bio-Rad Trans-Blot Turbo Transfer System set to constant 2.5 A (up to 25 V) for 7 minutes. Membranes were rinsed with 2 x 20 mL of 50 mM Tris·HCl pH 7.4, 150 mM NaCl (TBS) and blocked using 20 mL of TBS containing 5% BSA for 1 h at room temperature. After blocking, membranes were incubated with TBS, 0.1% Tween-20 (TBST) containing 5% BSA and 1.8 µg $mL^{-1}$ Alexa Fluor 680–conjugated streptavidin (Jackson Immunoresearch, 016-620-084) for 1 h at room temperature in the dark. Membranes were washed with 20 mL TBST for 5 min and imaged using a Bio-Rad ChemiDoc MP Imaging System with the Alexa 680 channel.

***Microbiota harvesting and processing***

All animal experiments were performed using procedures approved by the UC Irvine Institutional Animal Care and Use Committee in protocol AUP-25-071. Fresh fecal pellets were collected separately from 9–11-week-old, male C57BL/6NTac mice (Taconic Bioscience, B6-M) in individual 2-mL microcentrifuge tubes containing Lysing Matrix D (MP Biomedicals, 116913100) and 1 mL of deoxygenated PBS. Suspensions were homogenized by a FastPrep-24 5G bead beating grinder (MP Biomedical) using 4 m $sec^{-1}$ motor speed for 15 sec at room temperature. Samples were then transferred to a vinyl anerobic chamber (Type B, Coy Lab Products) maintained with an atmosphere of 90% $N_2$, 5% $CO_2$, and 5% $H_2$. Each sample corresponding to fecal homogenate from an individual mouse was processed separately as individual biological replicates. Suspensions were passed through a 35-micron strainer cap (Genessee, 28-155) into 15-mL culture tubes (Fisher Scientific, 07-000-212) to remove large particulate material. The filtered suspensions were then diluted with deoxygenated PBS to an OD600 of approximately 0.5.

***On-cell microbiota photocrosslinking and CuAAC labeling***

For analytical flow cytometry experiments, bacterial suspensions prepared from separate mice were pooled prior to photocrosslinking to measure the variability of labeling by each probe across an equivalent bacterial population. Bacterial suspensions were then treated with the indicated probe at the specified concentration in a total volume of 250 µL in non-treated 24-well culture plates (VWR, 10861-558). For flow sorting experiments, bacterial suspensions were maintained as individual biological replicates and incubated with 25 µM **2'-FL-1** in a total volume of 5 mL in non-treated 6-well tissue culture plates (VWR, 10861-554). Samples were gently mixed and incubated anaerobically for 30 min at RT in the dark. Samples were then irradiated under 365 nm light for 10 min on ice using a Spectrolinker XL-1000 UV crosslinker (Spectro-UV) with the samples positioned approximately 2 cm from the light source.

Following irradiation, analytical flow samples were transferred into 1.5-mL microcentrifuge tubes, and flow sorting samples were transferred to 15-mL conical tubes. Transferred samples were then centrifuged at 4,000 × *g* for 5 min, and the resulting cell pellets were washed twice with PBS with centrifugation at 4,000 × *g* for 5 min between washes. Washed cells were resuspended in 750 µL of PBS, and 250 µL of 4% formaldehyde diluted from 16% formaldehyde (Thermo Fisher Scientific, PI28908) was added for fixation. Samples were incubated for 30 min at room temperature and subsequently centrifuged at 8,000 × *g* for 5 min. The supernatant was removed, and the resulting cell pellets were washed twice with 1 mL of PBS.

CuAAC mix was prepared immediately before use by sequentially combining in order 2 parts of 50 mM CuSO4•5$H_2O$, 2.5 parts of 4 mM BTTAA, 2 parts of 100 mM sodium ascorbate, and 1 part of 10 mM AZDye 647 Azide Plus (Vector Laboratories, CCT-1482-25) followed by 93 parts of PBS, with mixing after each addition. Cell pellets were resuspended in CuAAC mix (50 µL for analytical samples and 500 µL for sorting samples). Samples were then incubated for 60 min at room temperature in the dark with shaking at 600 rpm and subsequently centrifuged at 8,000 × *g* for 5 min. The supernatant was removed, and cell pellets were washed once with 1% BSA in PBS (100 µL for analytical samples and 500 µL for sorting samples for all subsequent steps). Samples were then resuspended in 1% BSA in PBS, incubated for 5 min at room temperature with shaking at 600 rpm, and centrifuged at 8,000 × *g* for 5 min. Cell pellets were washed twice with PBS and stored in PBS at 4 °C until flow cytometric analysis.

##### *Analytical flow cytometry*

Analytical flow cytometry was performed using an Agilent NovoCyte three-laser flow cytometer (B5-YG5-R3). Samples were resuspended in 100 µL of filtered flow buffer (1% BSA in PBS) and transferred to a non-treated 96-well, V-bottom plate (Fisher Scientific, 07-200-698). Cells were then treated with 100 µL of PBS containing 5 µM SYTO 21 (ex/em 494/517 nm; Thermo Fisher Scientific, S7572) immediately prior to analysis. Data acquisition was performed with an FSC-H threshold of 10,000 and continued until either 40,000 events within the SYTO $21^+$ population were collected or 90 µL of sample had been analyzed. Gates were drawn using a no-probe sample as a negative control. Bacteria were gated as SYTO $21^+$ based on the B525-A and SSC-A channels to exclude debris. Labeled cells were subsequently gated as AZDye $647^+$ based on the R660-A and SSC-A channels. The flow cytometric gating strategy and representative samples are shown in **Figure S2**. Values are reported as percentage of AZDye $647^+$ cells within the SYTO $21^+$ population.

##### *Fluorescence-activated cell sorting*

Flow sorting samples were analyzed and sorted on a four-laser Sony SH800 Cell Sorter equipped with 70-µm sorting chips. Samples were resuspended in 15 mL of filtered flow buffer (1% BSA in PBS). Events corresponding to bacteria were first gated based on the FSC-A and BSC-A channels. Probe labeling was then measured using the APC-A and BSC-A channels, and the sorting gate for labeled cells was drawn using a no-probe sample as a negative control. The flow cytometric gating strategy and representative samples are shown in **Figure S3**. For each sample, 6,000,000 events were collected separately for the total bacteria ("all") and AZDye $647^+$ populations ("labeled"). Sorted cells were then centrifuged at 14,000 x *g* for 10 min. The supernatant was removed, leaving approximately 1 mL of residual volume, and the concentrated cell suspension was transferred to PowerBead tubes containing 0.1 mm glass beads (Qiagen 13118-50). Samples were stored at –80 °C and shipped on dry ice to the Weill Cornell Microbiome Core for further processing.

##### *DNA extraction and library preparation*

DNA extraction was performed using a Maxwell RSC PureFood GMO and Authentication Kit according to the manufacturer's protocol (Promega, AS1600). To each sample, 1 mL of CTAB buffer and 20 µL of RNAse A solution was added, and the tube was vortexed for 10 s using a Vortex Genie 2. Samples were then incubated at 95 °C with shaking at 1500 rpm for 5 min using an Eppendorf Thermomixer F2.0. Following incubation, tubes were secured horizontally using a microtube attachment (SI-H524) on a Vortex Genie 2 and vortexed at high speed for 20 min. After lysis, samples were centrifuged at 12,700 rpm at 4 °C for 2 x 10 min to reduce residual foam. Any remaining foam was carefully removed by pipetting to prevent obstruction of the robotic instrument tips.

Sample tubes were transferred to a Promega MaxPrep Liquid Handler. The instrument was loaded with proteinase K, lysis buffer, elution buffer, pipette tips, 96-sample deep well plate, and Promega Maxwell RSC 48 plunger tips according to the manufacturer's protocol. The instrument was programmed to transfer 300 µL of sample lysate to a Maxwell RSC 48 extraction cartridge. Following automated sample preparation, the cartridge was transferred to a Maxwell RSC 48 instrument for DNA extraction and elution. DNA was eluted in 100 µL of elution buffer and transferred to a standard 96-well plate. DNA concentrations were quantified using the Quant-iT dsDNA High Sensitivity Assay Kit (Thermo Fisher Scientific, Q33120) using a GloMax plate reader with a Greiner Bio-One CELLSTAR microplate (Promega, 655087).

Metagenomic shotgun sequencing libraries were prepared using the Nextera XT DNA Library Prep Kit (Illumina) according to the manufacturer's instructions. Sequencing libraries were purified using AMPure XP magnetic beads (Beckman Coulter) according to the manufacturer's instructions. Library size and quality were verified using a Labchip GX Touch HT instrument (Revvity) with DNA 1K Reagent Kit (Revvity, CLS760673). Library concentrations were quantified using the Quant-iT dsDNA High Sensitivity Assay Kit as described above. Library molarity was calculated based on the measured DNA concentrations and library peak sizes. Libraries were normalized to 2 nM using a NIMBUS instrument (Hamilton Microlab). Equal volumes of the normalized libraries were then pooled into a 1.5 mL DNA LoBind tube (Eppendorf, 022431021).

##### *Sequencing and data processing*

Pooled libraries were sequenced on an Illumina NovaSeq X instrument at a loading concentration of 0.13 nM with 5% PhiX spike-in using 2x150 bp paired-end sequencing. Raw sequencing read files are available in BioProject PRJNA1545266. Raw sequencing reads were processed using KneadData (v. 0.12.4, Huttenhower lab, https://huttenhower.sph.harvard.edu/kneaddata/) for quality-based filtering and adapter trimming via Trimmomatic (v. 0.41).[1] Host reads were identified by Bowtie2 (v. 2.5.5)[2] using the RefSeq GRCm39 mouse reference genome and removed. Taxonomic profiles were generated using MetaPhlAn 4 (v. 4.2.5, marker database mpa_vJan26_CHOCOPhlAnSGB_202605)[3] using default parameters except skipping unclassified estimation, and the read-to-marker alignments were retained (--mapout). Because the reads_map analysis type is currently unavailable in v. 4.2.5, per-species read counts were obtained by tallying the per-read species assignments reported by reads_map in MetaPhlAn 4 (v. 4.1.1) run against the same marker database. Reads aligning to markers of more than one species were excluded, yielding 792,883-1,747,064 species-assigned reads per sample across 2,368 species-level taxa. Raw assigned read values are provided in **Table S4**.

##### *Enrichment analysis*

To remove low-abundance taxa, the initial read count dataset was filtered to species representing ≥0.05% of species-assigned reads in at least four of the eight samples, yielding 105 species. Each count was increased by 0.5 reads to permit logarithmic transformation of zero values, and relative counts were calculated as the fraction of total species-assigned reads for that sample, with the total values derived from unadjusted read counts across all 2,368 species (see **Table S5**). Log2-transformed values for AzDye $647^+$ ("labeled") and total ("all") samples from the same mouse were treated as pairs ($n = 4$ mice), with fold changes expressed as labeled relative to all. Pairs were analyzed using a paired $t$ test with the two-stage Benjamini, Krieger, and Yekutieli procedure[4] to control the false discovery rate (Prism 11, v. 11.0.2). A $q$ value of 0.1 was used as the significance cut-off.

##### *Cladogram visualization*

Enriched or depleted species were visualized by a cladogram produced by iTOL v. 7.6[5] based on the maximum-likelihood tree of all species-level genome bins (SGBs) that is distributed with the MetaPhlAn marker database listed above. The 105 species-level taxa were subsetted to those that mapped to a single SGB for visualization only (101 species-level taxa; excluding *Lachnospiraceae bacterium*, *Clostridia bacterium*, *Eubacteriaceae bacterium*, and *Oscillospiraceae bacterium*), and the reference tree was pruned to these 101 SGBs using the keep.tip function in ape (v. 5.8.1) in R (v. 4.5.3) and uploaded into iTOL. Branch lengths were set to "Ignore" and scaling factors were set to 0.5x to produce the cladogram structure. Individual tips were annotated using the absolute value of log2(fold change) calculated in Prism 11 from the previous step. Symbols were semi-proportionally sized based on the following formula: $3 + 23 \times |\log2(FC)|/3.6709$, where 3 was used as a floor to prevent symbols from vanishing and 3.6709 is the largest $|\log2(FC)|$ value. Symbols were colored only if $|\log2(FC)| > 1$ and $q < 0.1$, with colors denoting the direction of the enrichment or depletion. All statistically significant SGBs fulfilled these criteria and were colored on the cladogram. Bacillota and Bacteroidota phyla were annotated as a color strip based on the clade_name value of each tip, and all remaining phyla were left unannotated.

##### *Data analysis and statistics*

Gel images were processed and quantified using Bio-Rad Image Lab software (version 6.1). All raw gel images are provided in **Figures S4-S6**. All graphical representations and quantification for flow cytometric data were generated using FlowJo 10 (ver. 10.10.0), FlowJo 11 (ver. 11.1.0, Becton Dickinson & Company), and NovoExpress software (ver. 1.5.0, Agilent). Unless otherwise noted, graphical representations and statistical analyses of quantitative data were generated in Prism (ver. 10.6.0, GraphPad). All graphed values are provided in **Table S6**.

For gel quantification, densitometric values for lectin labeling were normalized to total protein loading and compared using an ordinary one-way ANOVA with Tukey's multiple comparisons post-hoc test. For comparisons between ratios of Bacillota and Bacteroidota or between normalized read counts of Clostridiaceae, paired ratio t-tests were employed. Statistical enrichment or depletion of taxa within a phylum was quantified by two-tailed Fisher's exact tests. For read count comparisons of all species-level taxa between all and labeled samples, log-transformed, normalized read counts as described above were compared using a paired $t$ test with the two-stage Benjamini, Krieger, and Yekutieli procedure to control the false discovery rate. Where possible, exact statistical test values are reported.

#### NMR spectra

$^{1}H$ and $^{13}C\{^{1}H\}$ spectra for all synthesized compounds are provided on the following pages.

$^{1}$H NMR (500 MHz, $CDCl_3$) spectrum of compound **S1**

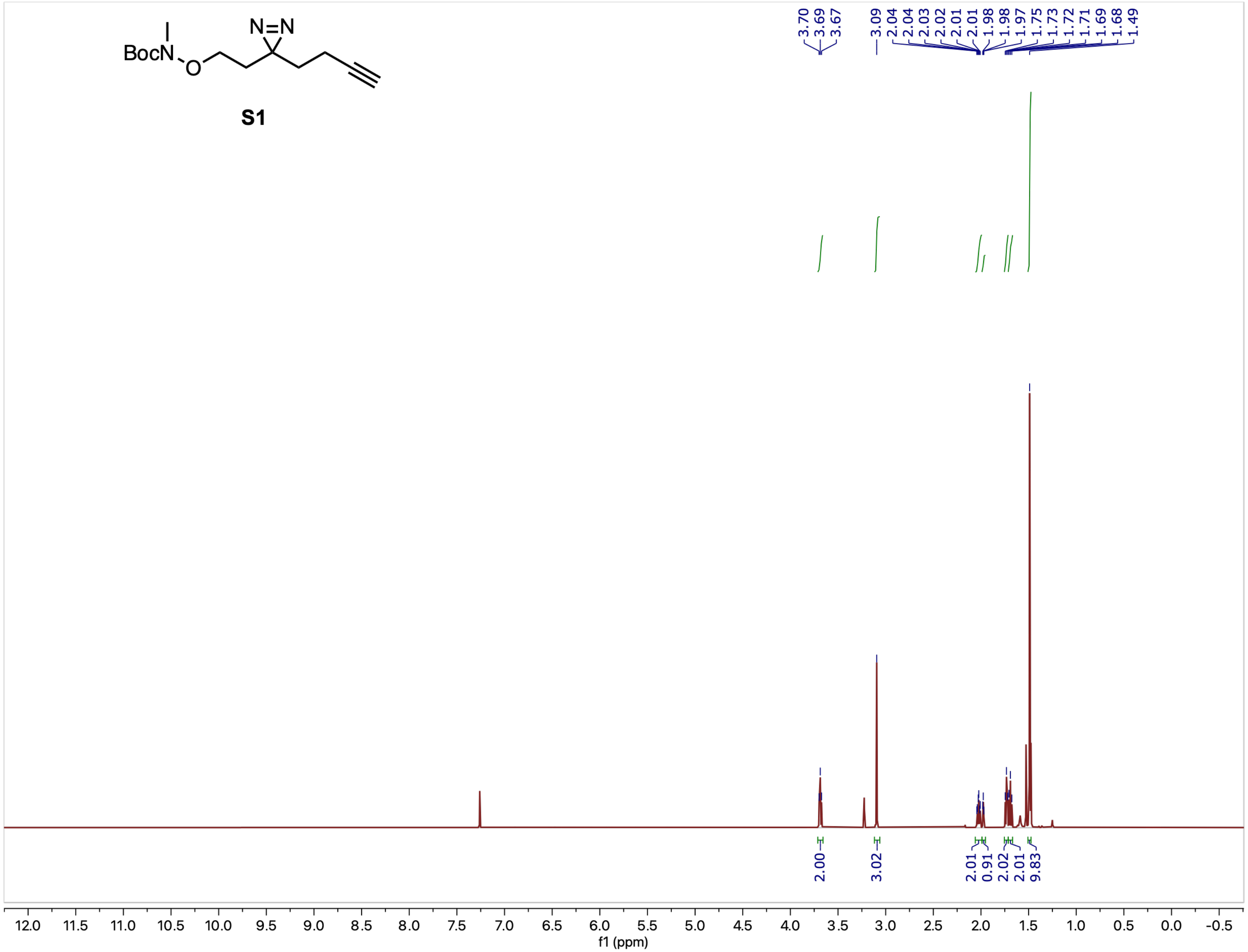

$^{13}C$ NMR (126 MHz, $CDCl_3$) spectrum of compound **S1**

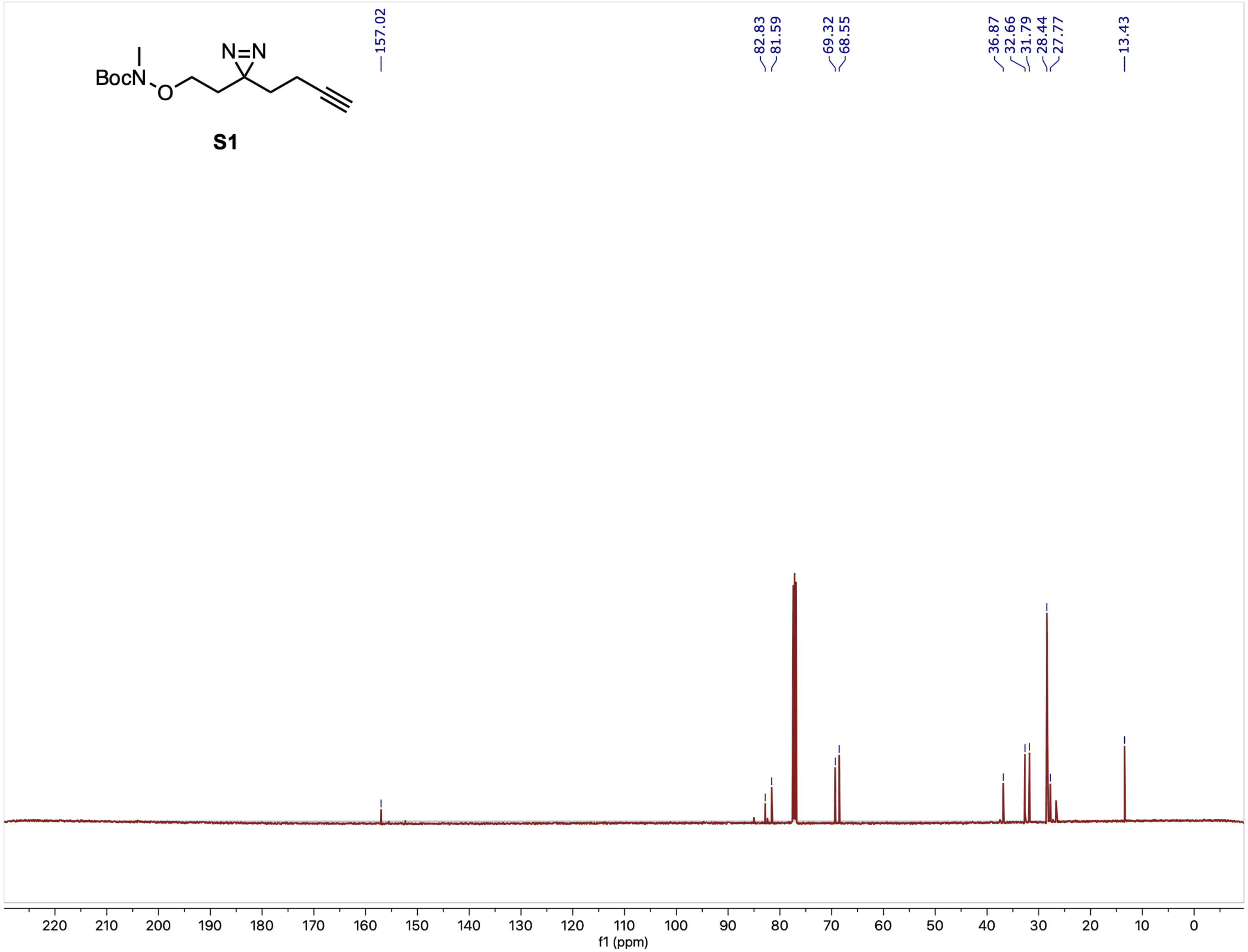

$^1H$ NMR (500 MHz, DMSO-$d_6$) spectrum of compound **1**

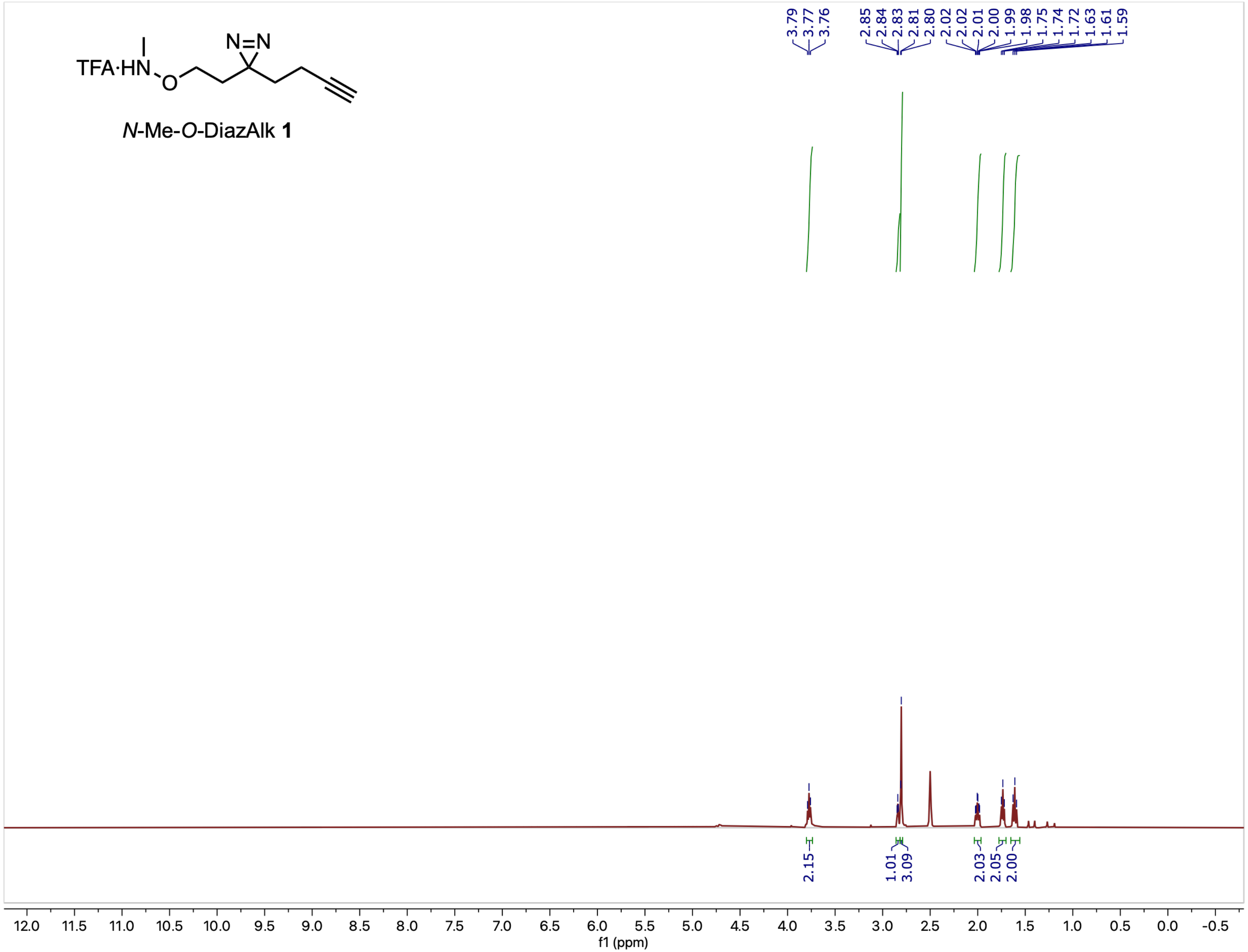

$^{13}C$ NMR (126 MHz, DMSO-$d_6$) spectrum of compound **1**

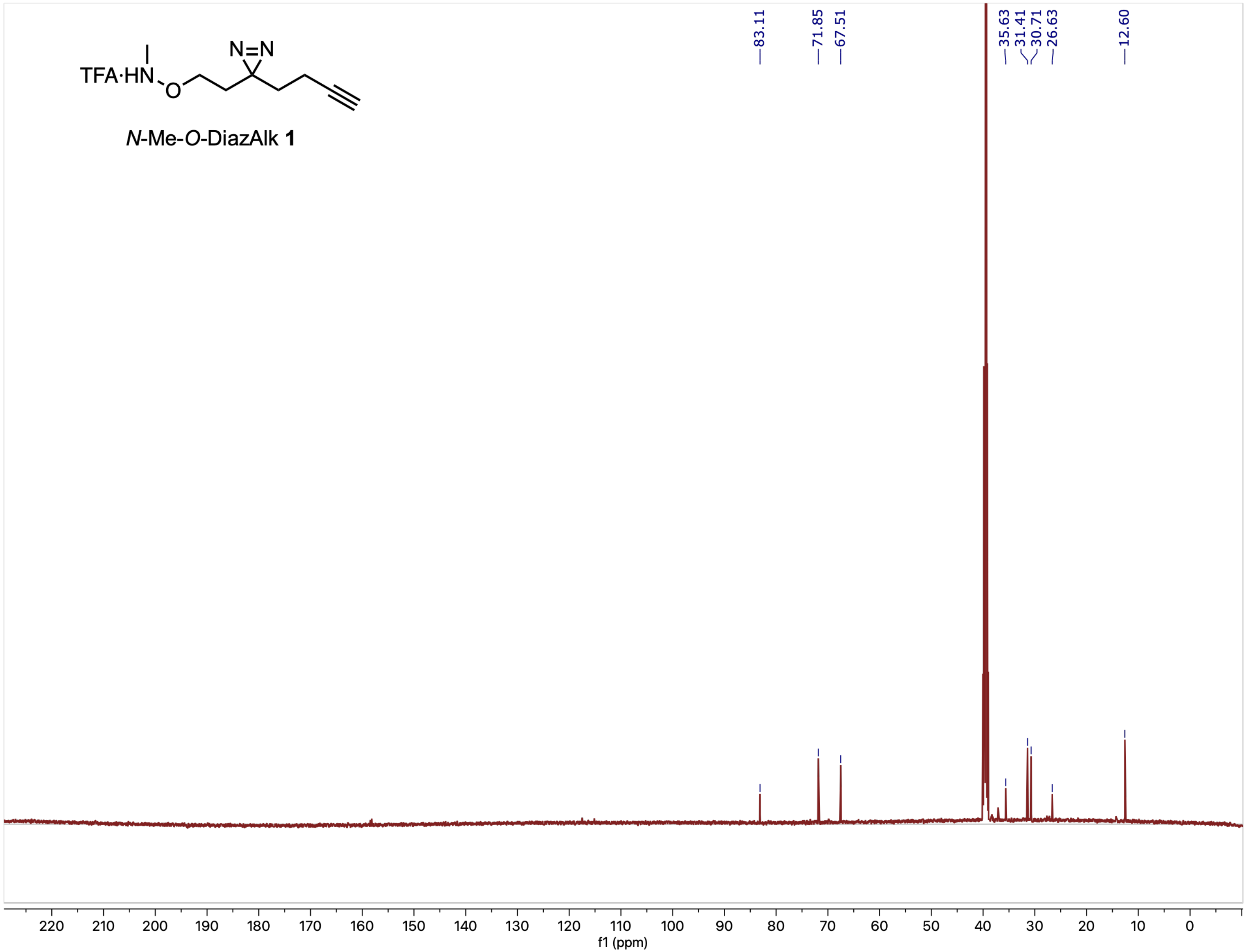

[1]H NMR (500 MHz, $CDCl_3$) spectrum of compound **S2**

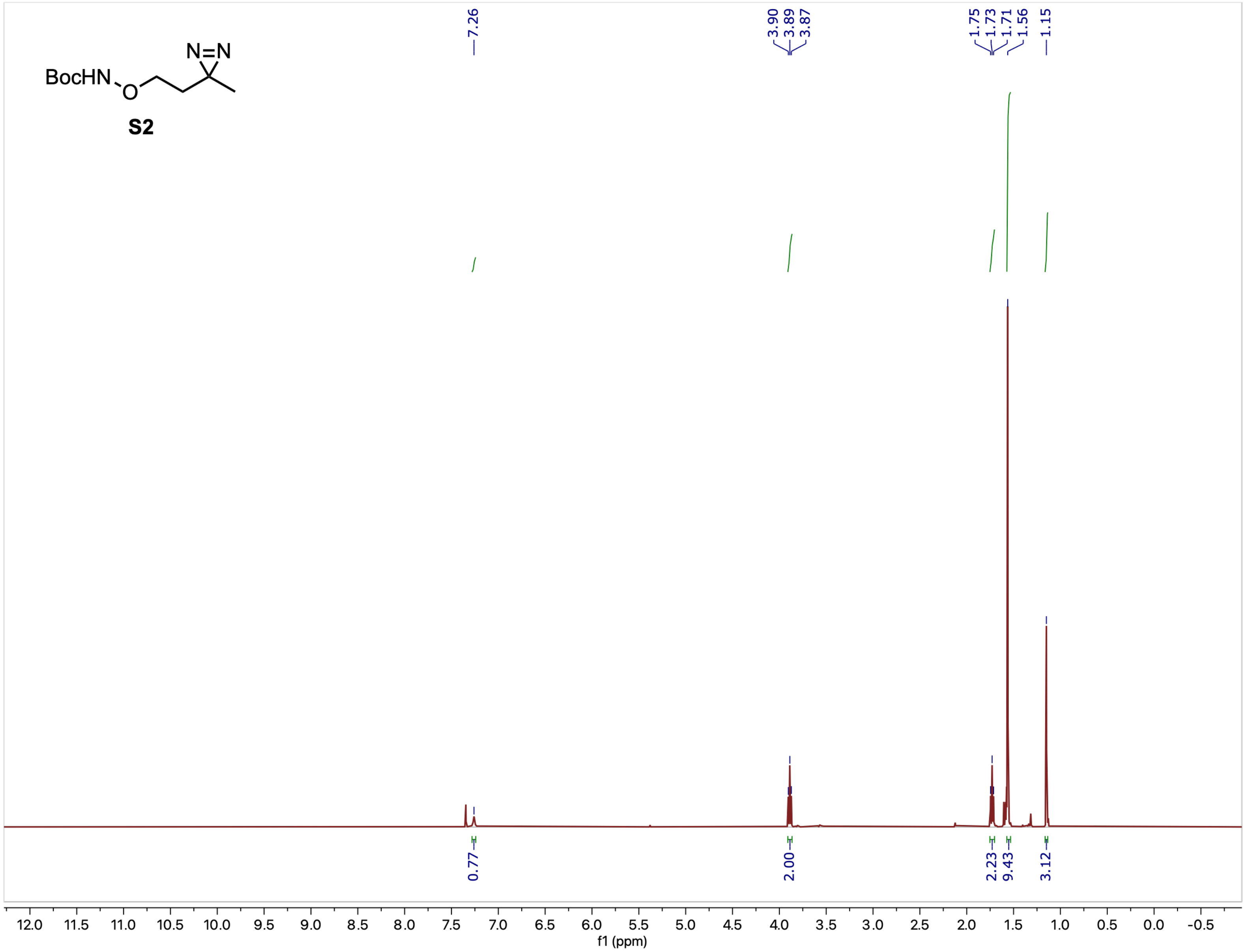

$^{13}C$ NMR (126 MHz, $CDCl_3$) spectrum of compound **S2**

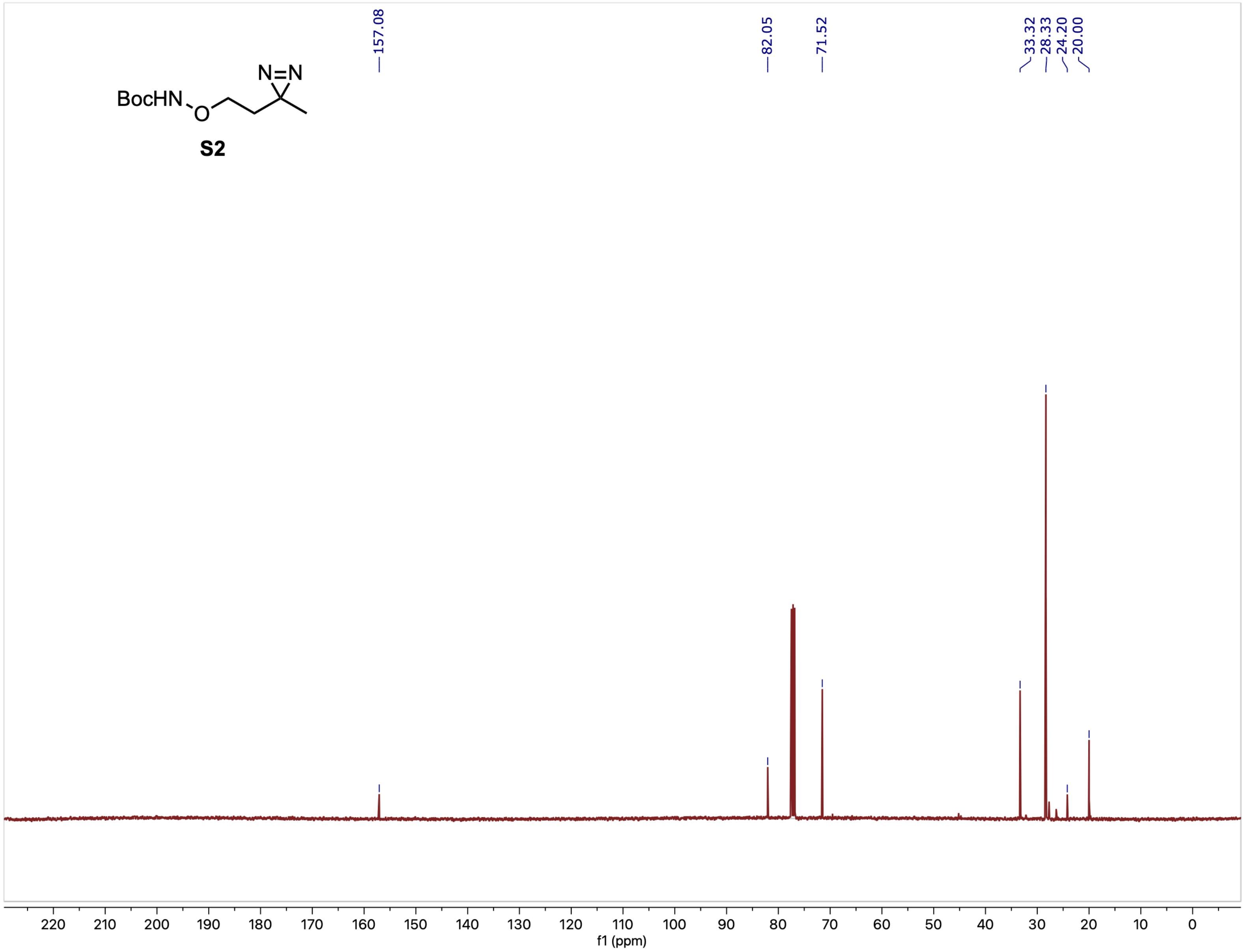

$^1$H NMR (500 MHz, $CDCl_3$) spectrum of compound **Boc-2**

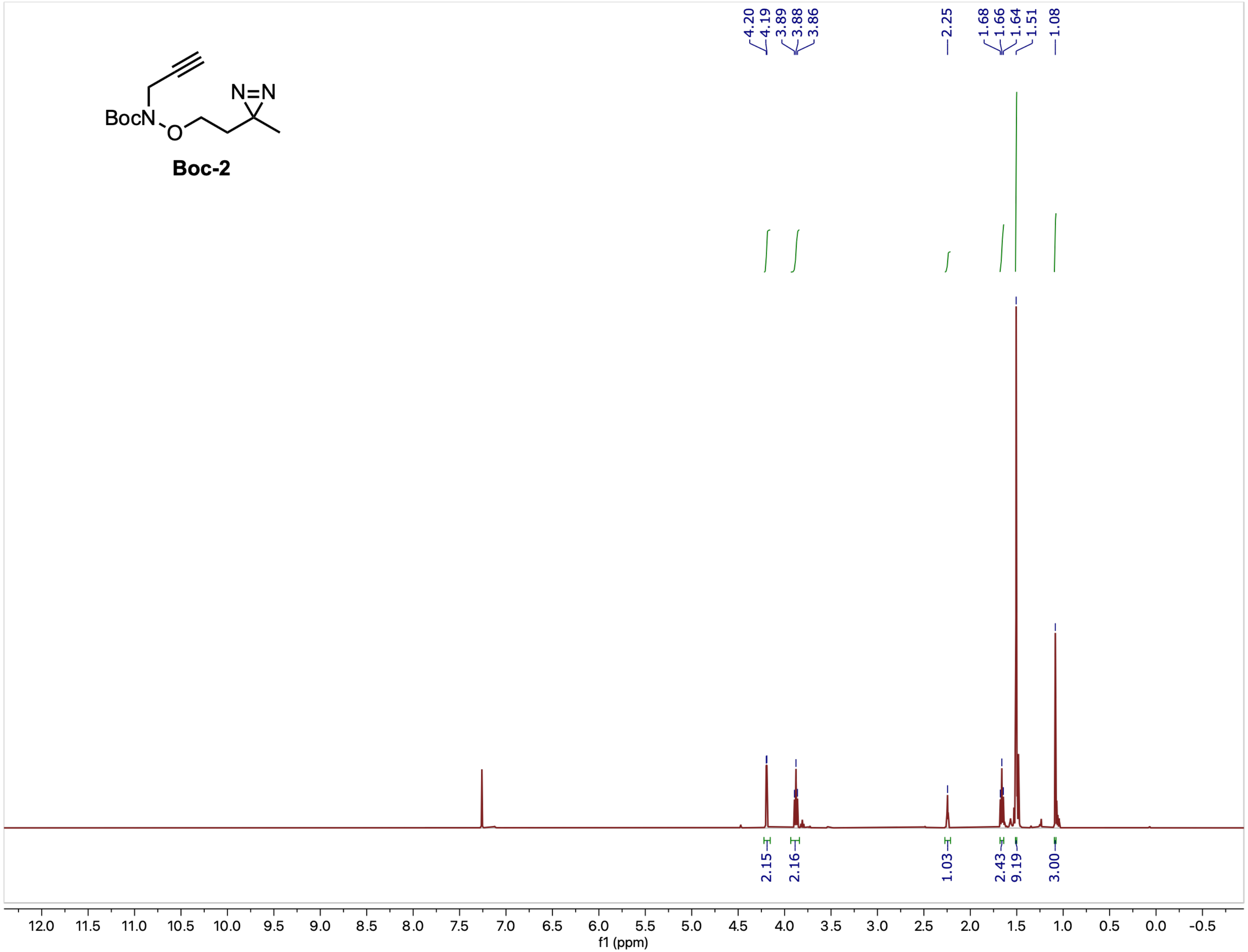

$^{13}C$ NMR (126 MHz, $CDCl_3$) spectrum of compound **Boc-2**

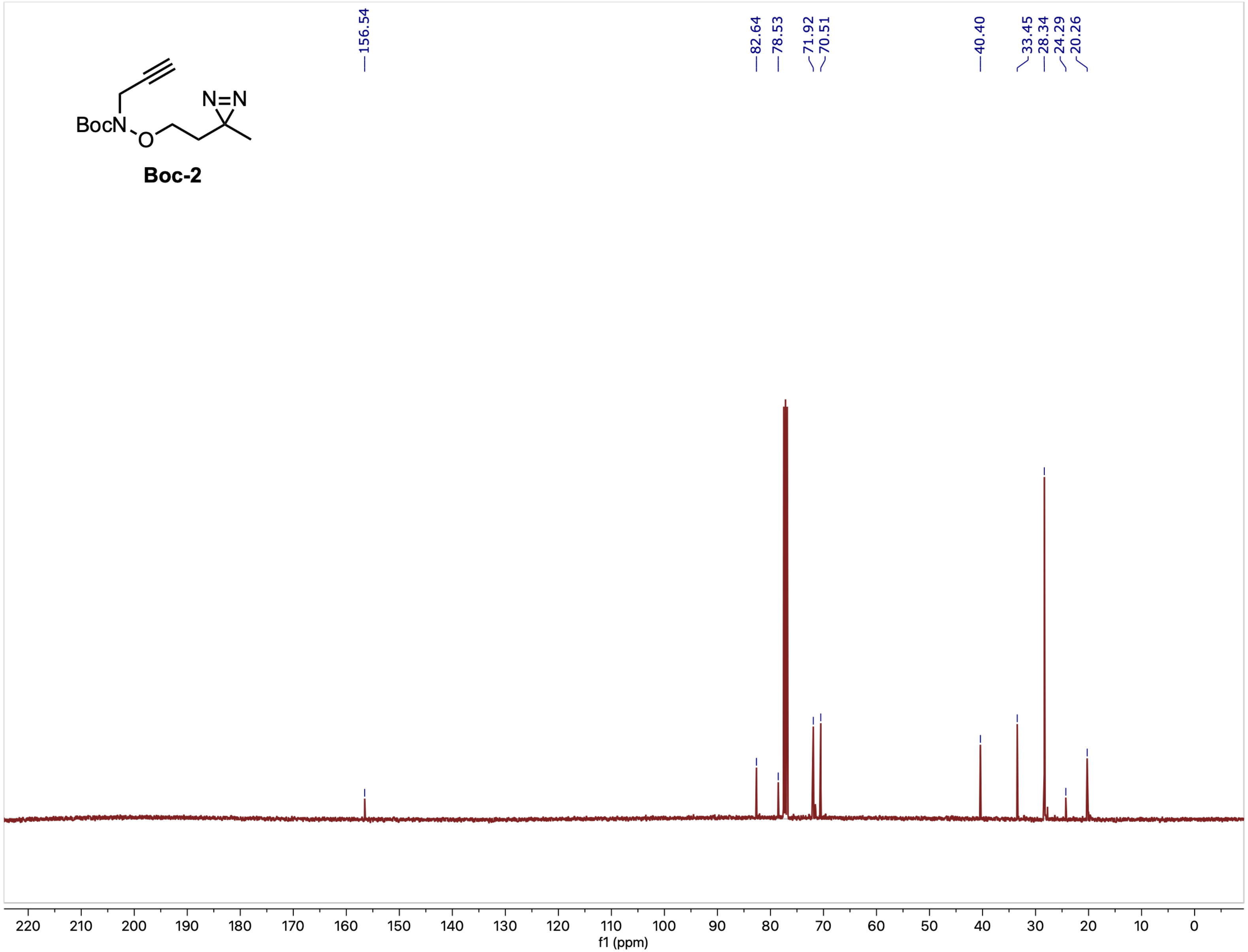

[1]H NMR (500 MHz, $CDCl_3$) spectrum of compound **S3**

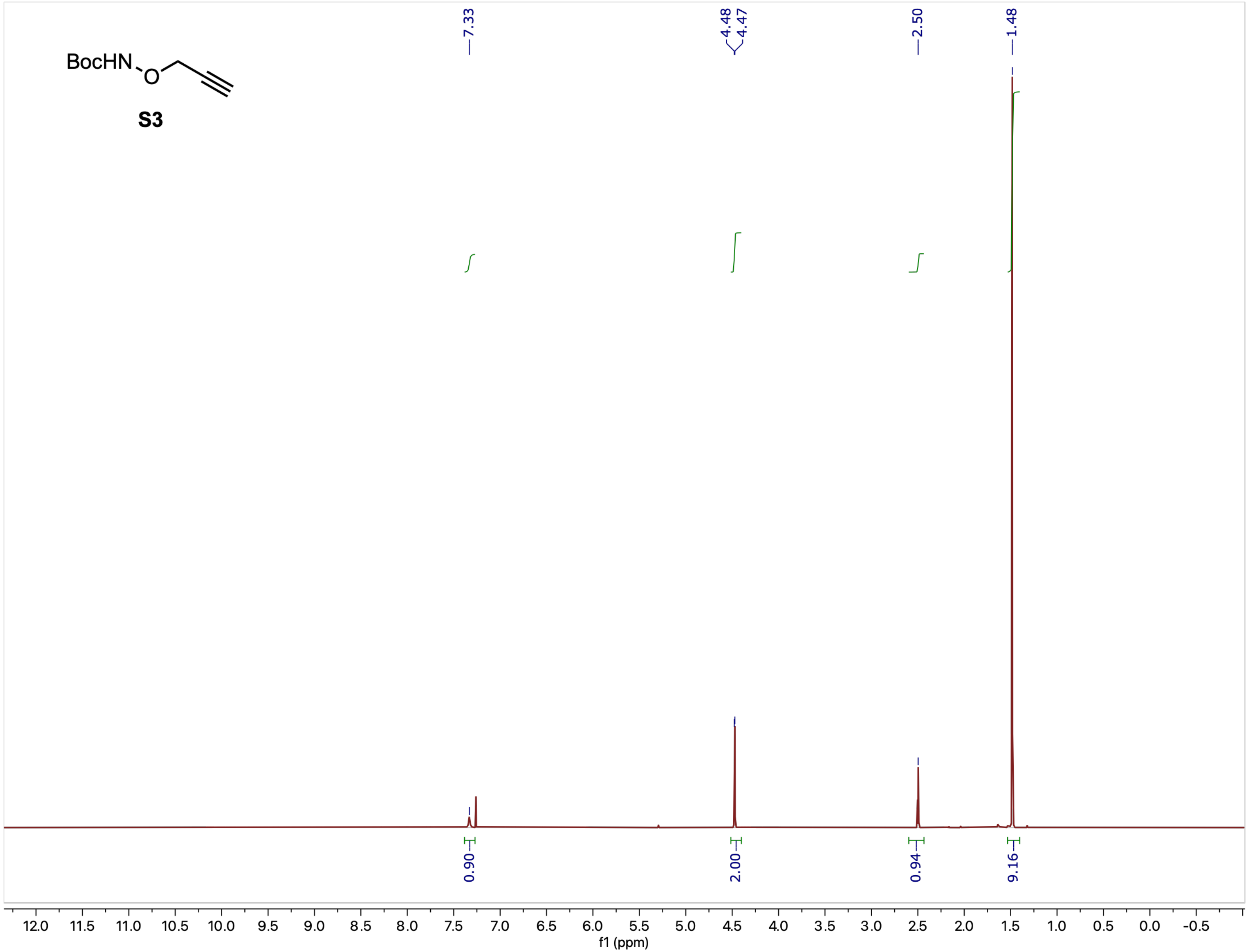

$^{13}$C NMR (126 MHz, $CDCl_3$) spectrum of compound **S3**

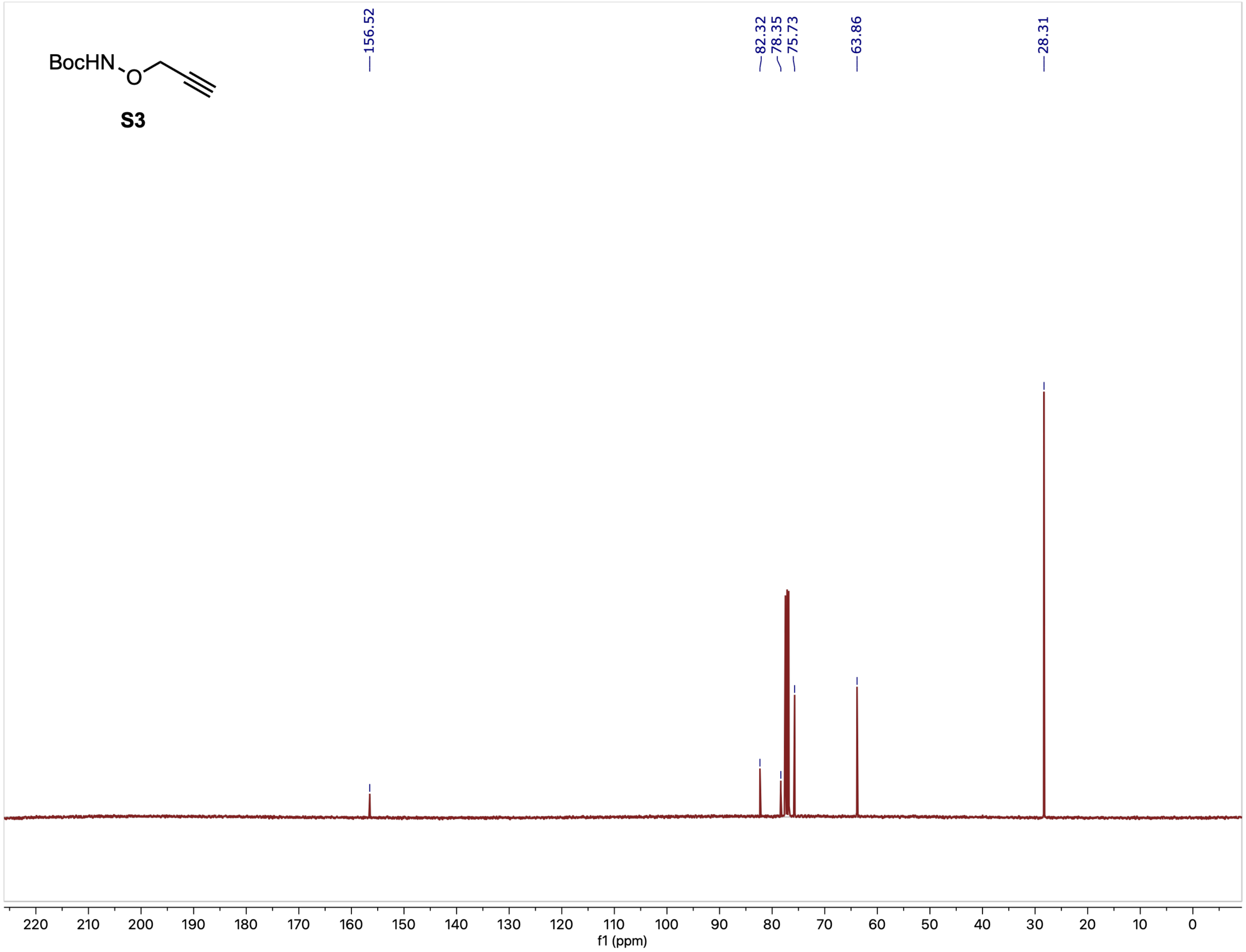

[1]H NMR (500 MHz, $CDCl_3$) spectrum of compound **Boc-3**

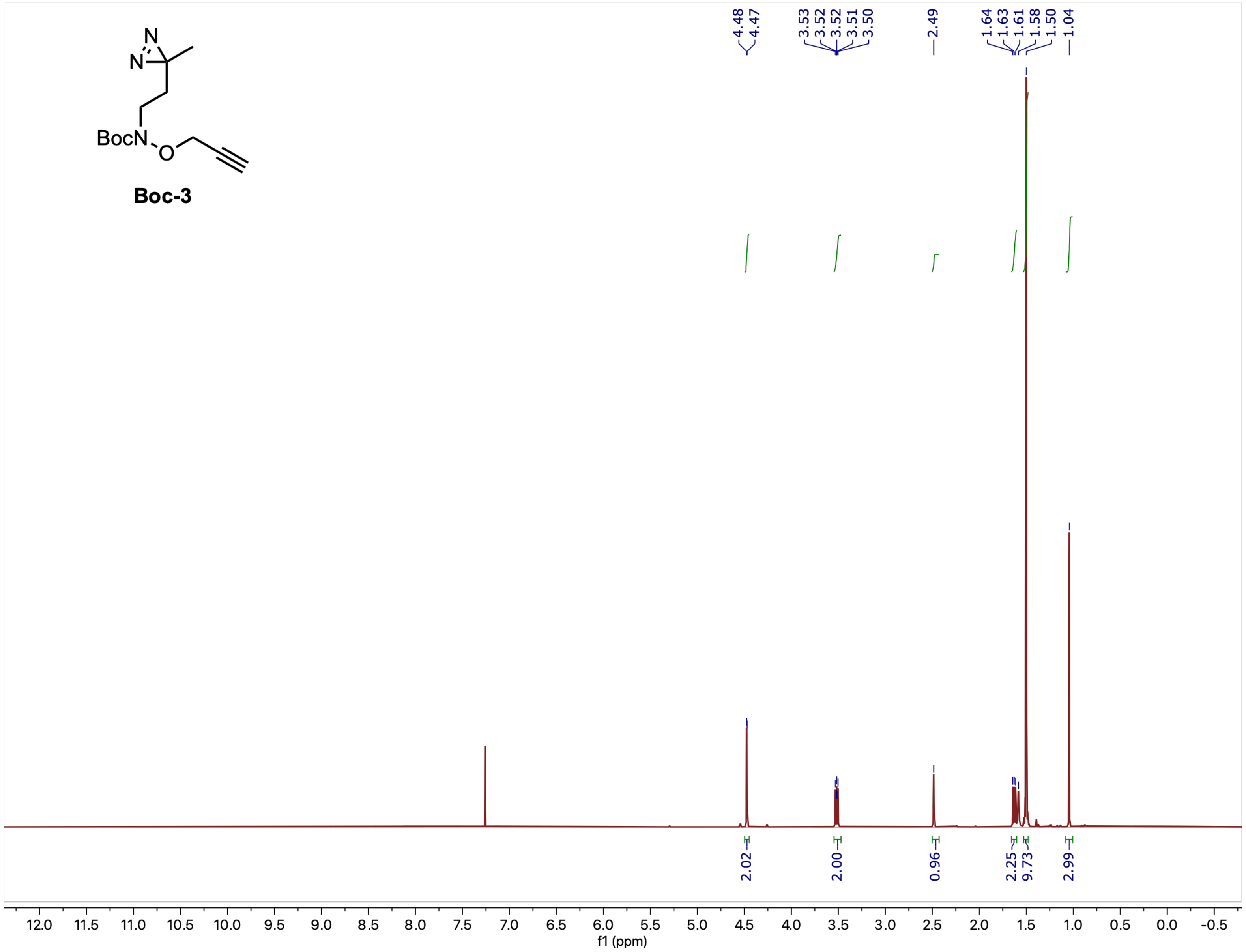

$^{13}C$ NMR (126 MHz, $CDCl_3$) spectrum of compound **Boc-3**

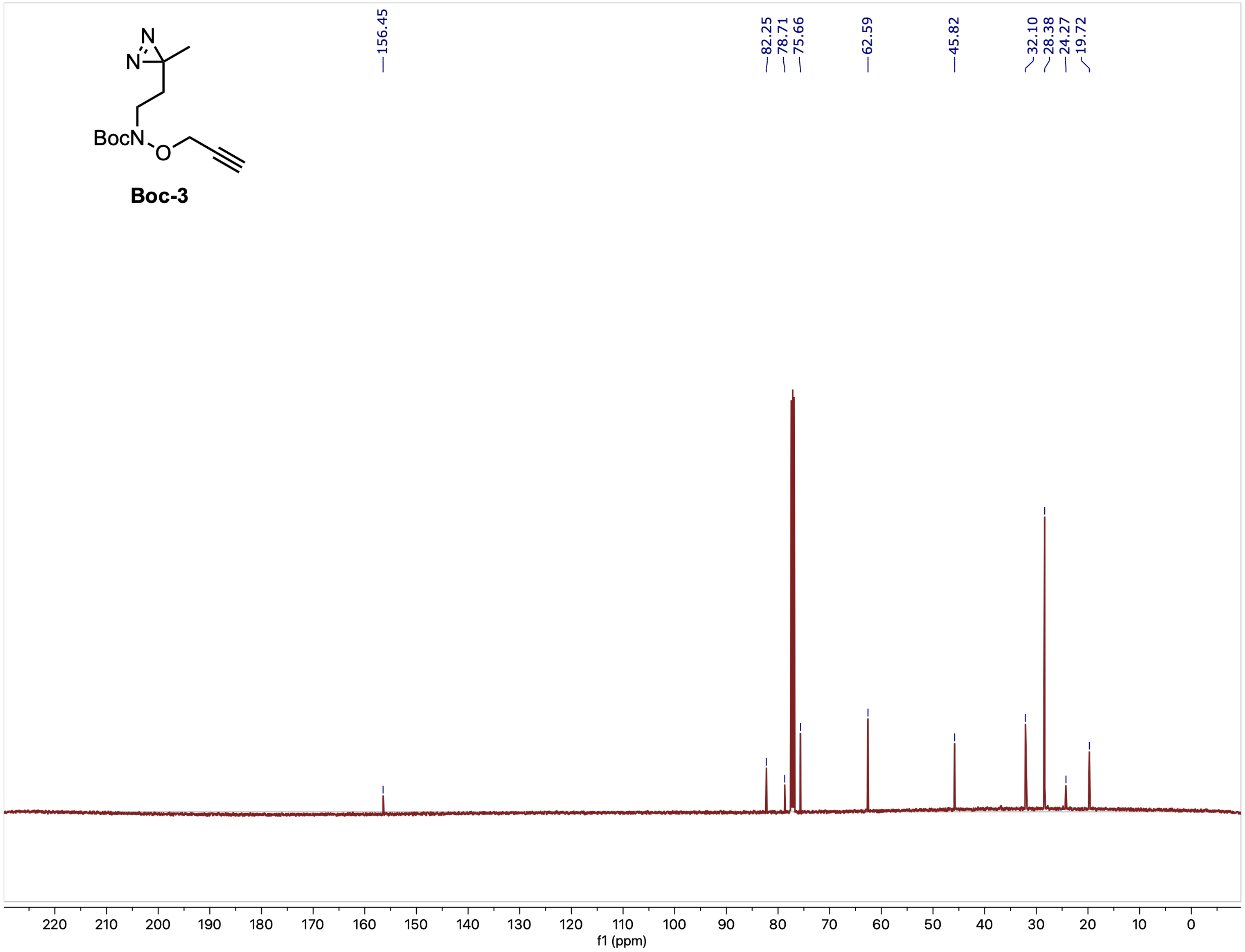

[1]H NMR (500 MHz, $CD_3OD$) spectrum of compound **Lac-1**

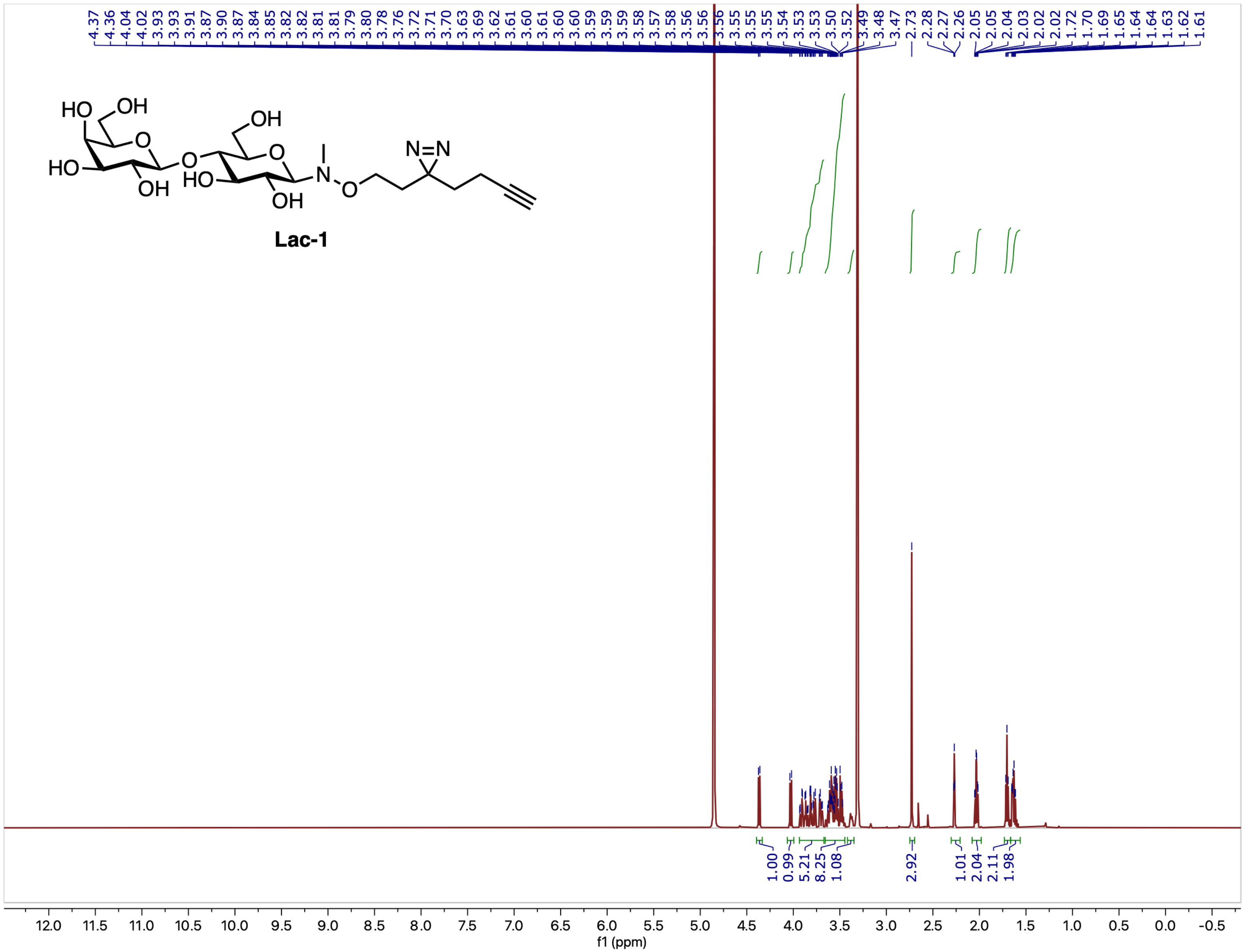

$^{13}$C NMR (126 MHz, $CD_3OD$) spectrum of compound **Lac-1**

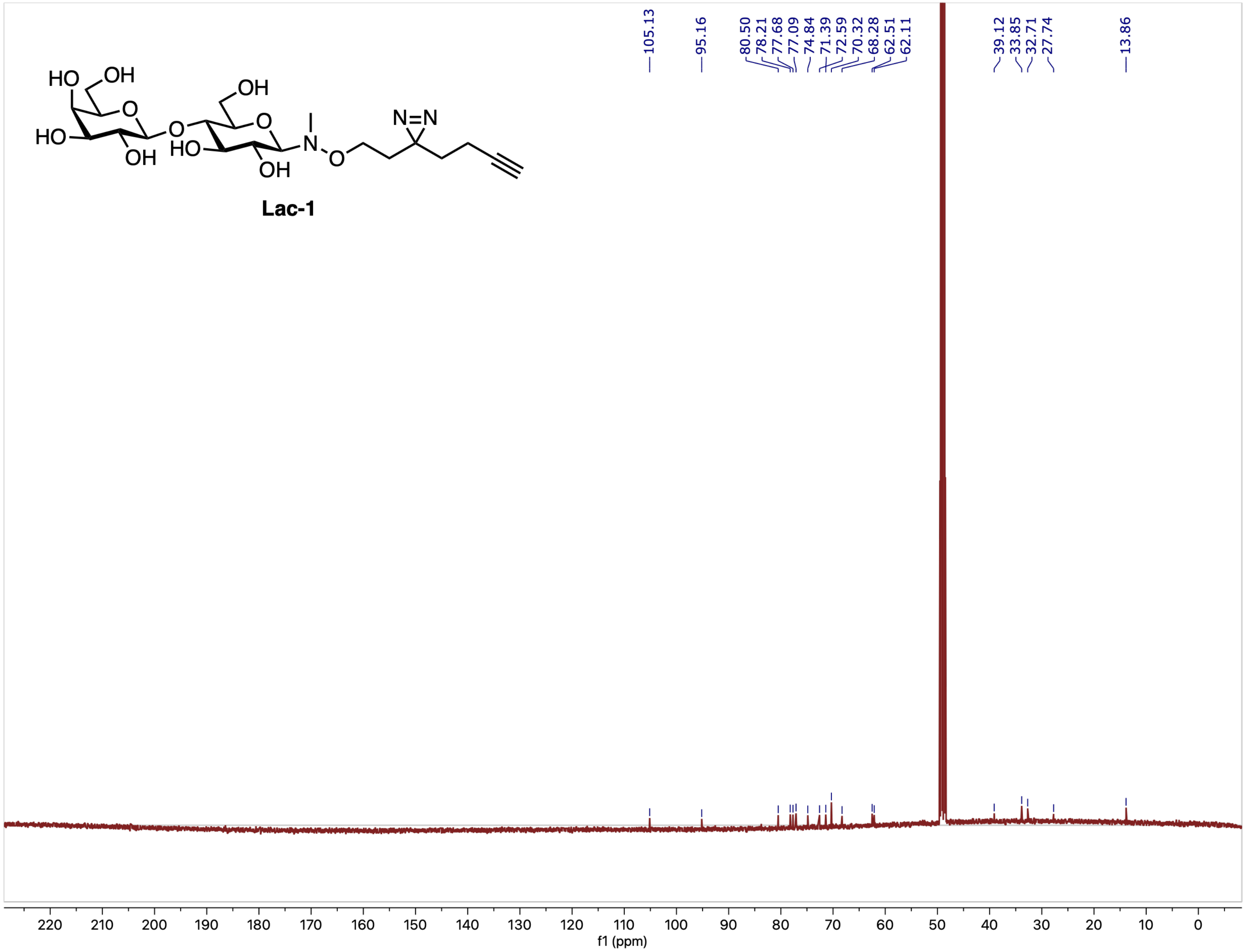

$^1$H NMR (500 MHz, $CD_3OD$) spectrum of compound **Lac-2**

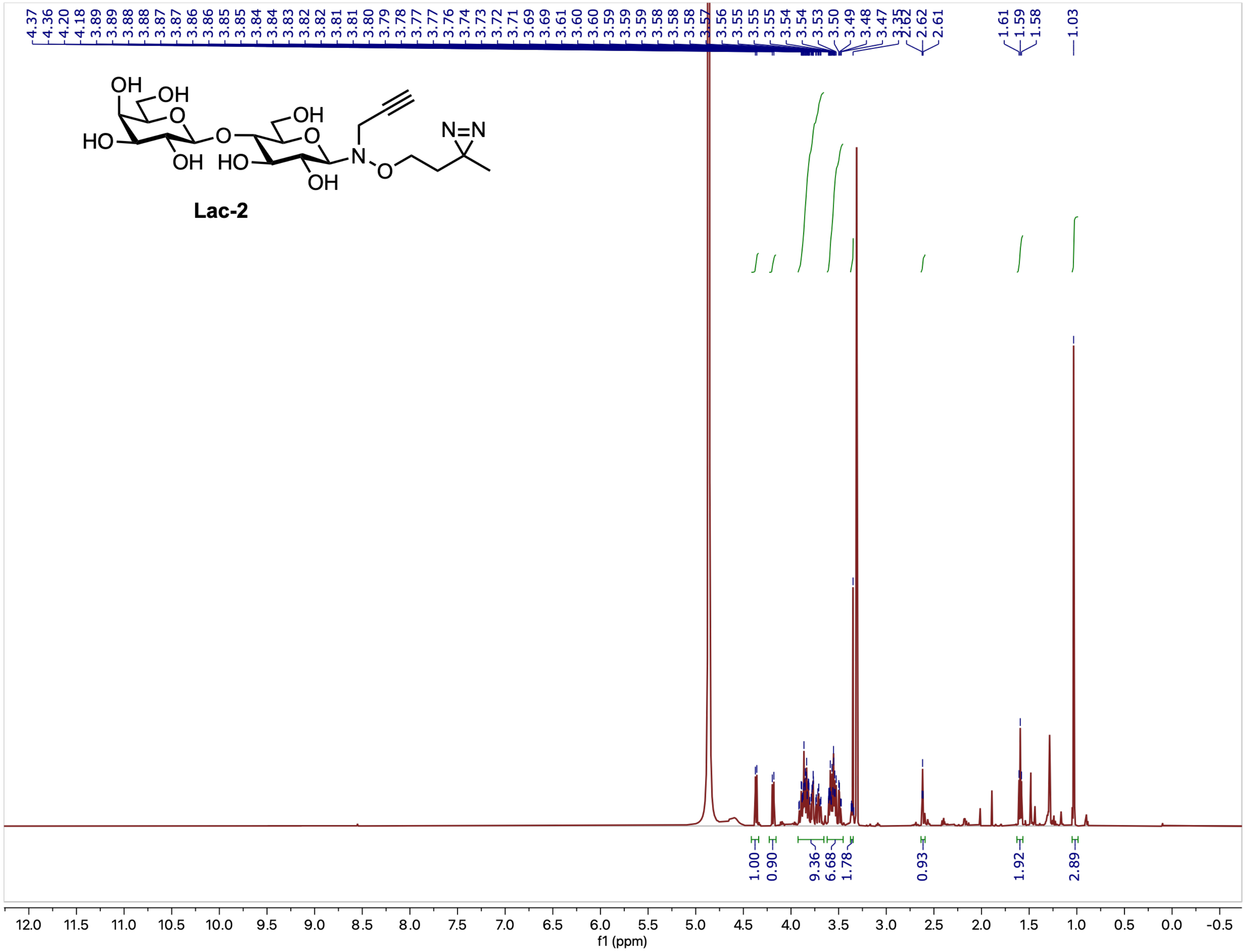

$^{13}C$ NMR (126 MHz, $CD_3OD$) spectrum of compound **Lac-2**

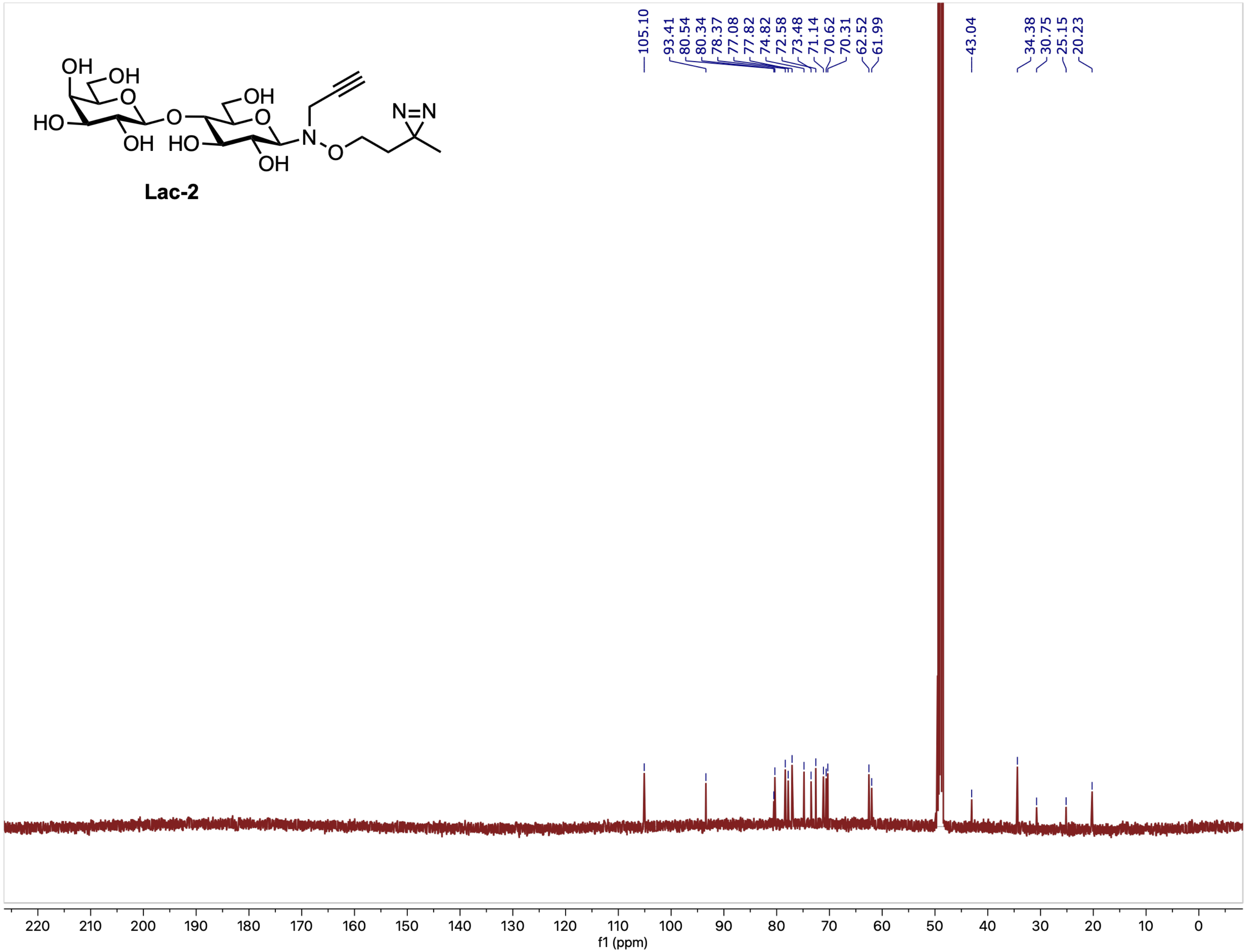

$^{1}$H NMR (500 MHz, $CD_3OD$) spectrum of compound **Lac-3**

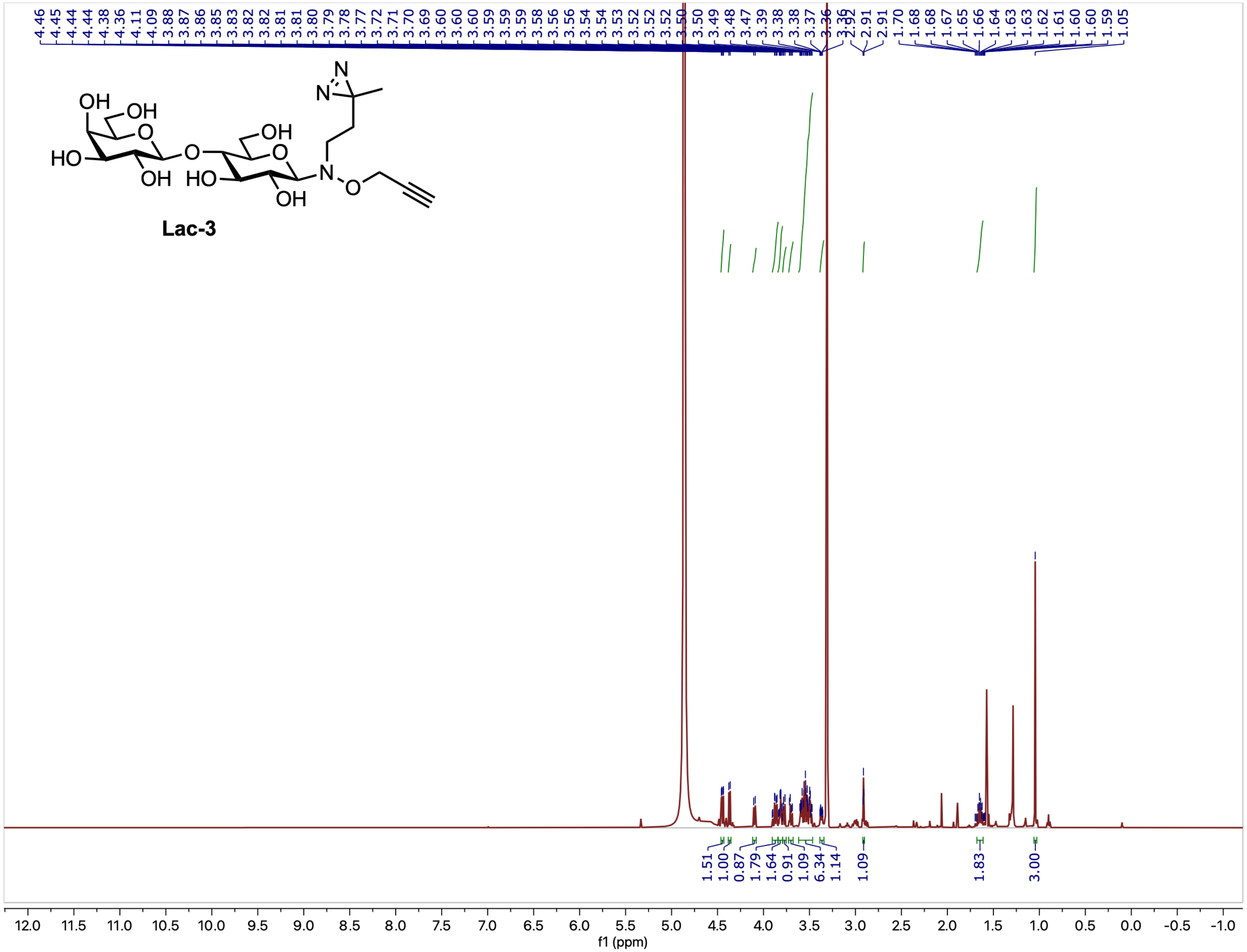

$^{13}C$ NMR (126 MHz, $CD_3OD$) spectrum of compound **Lac-3**

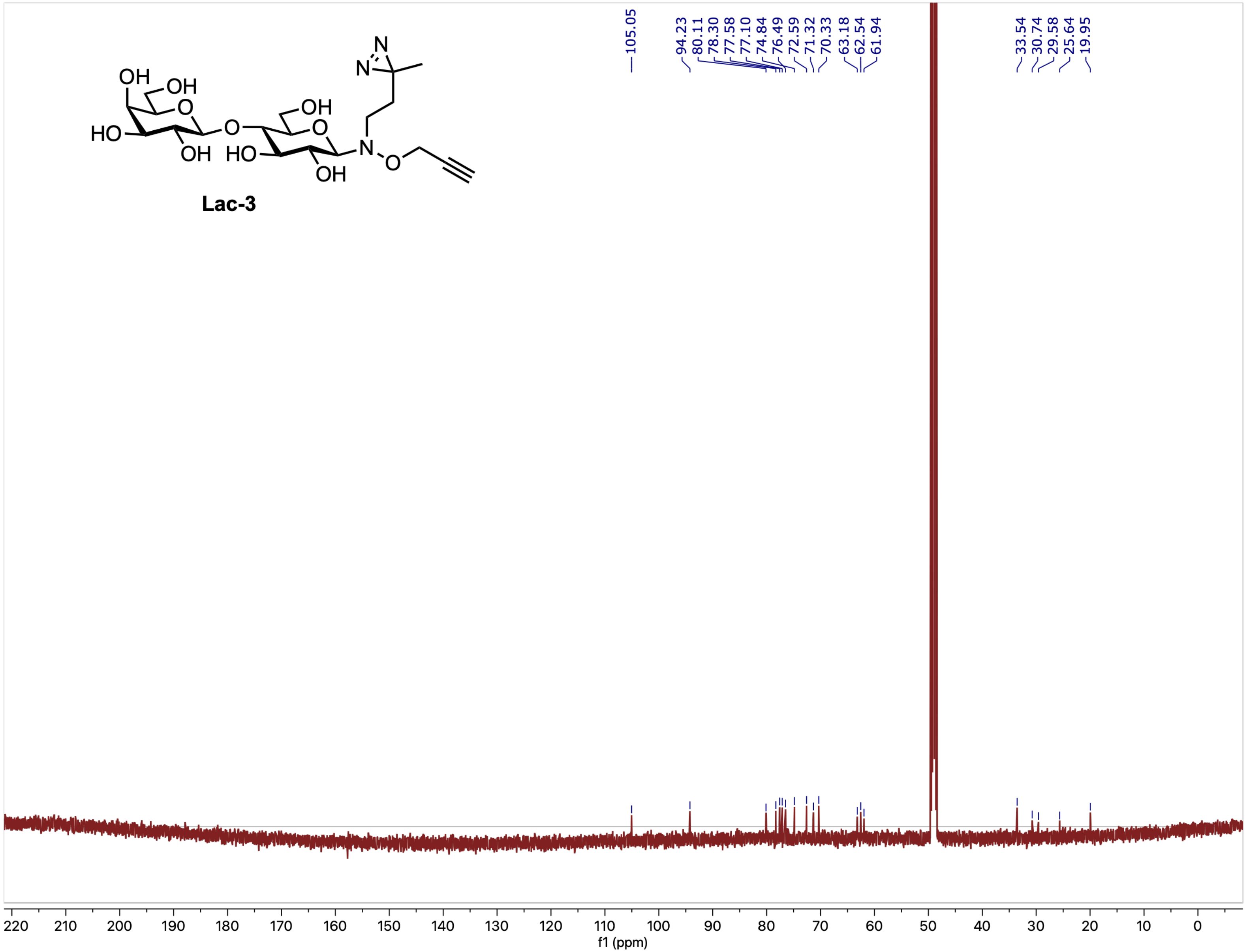

[1]H NMR (500 MHz, $CD_3OD$) spectrum of compound **2'-FL-1**

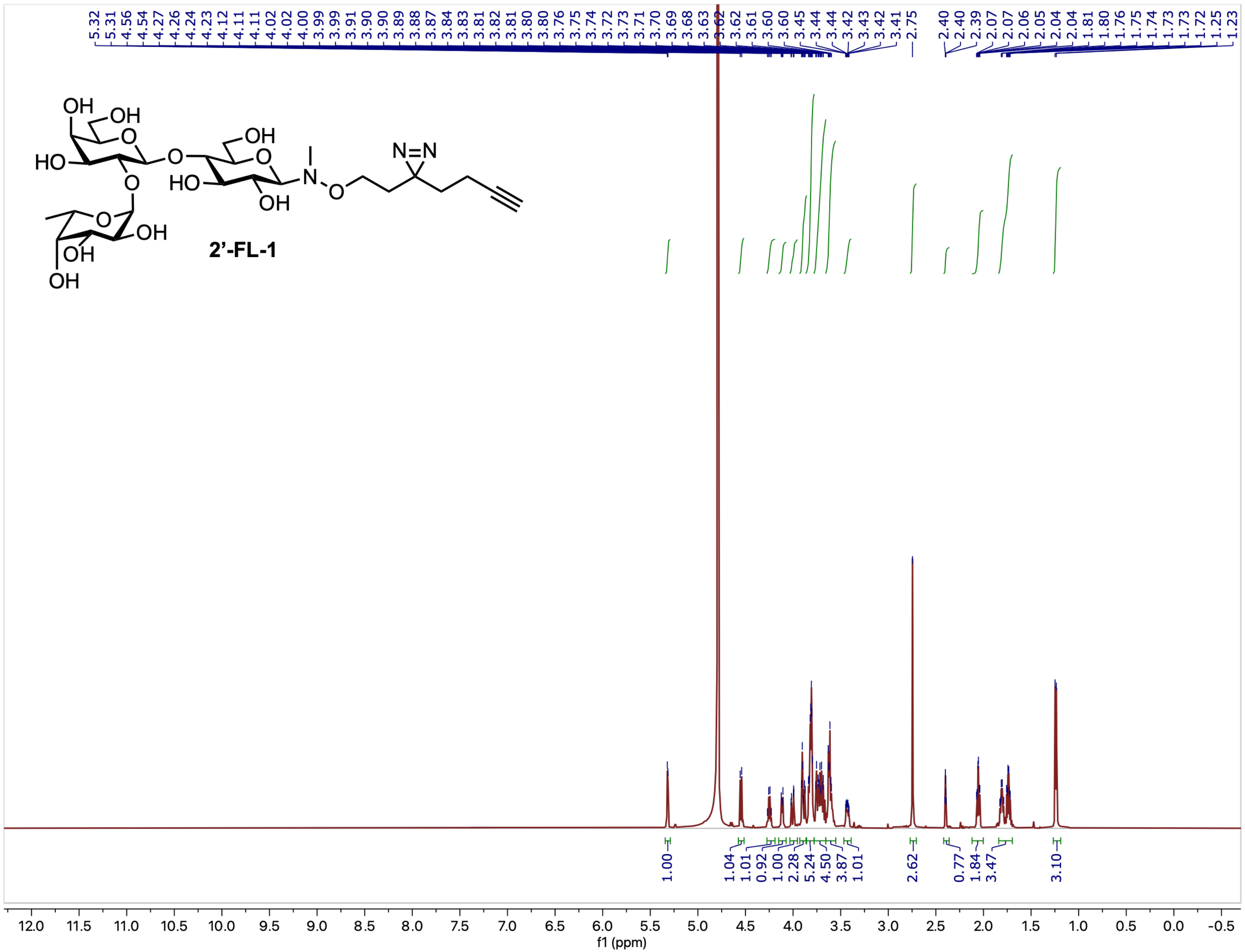

$^{13}C$ NMR (126 MHz, $CD_3OD$) spectrum of compound **2’-FL-1**

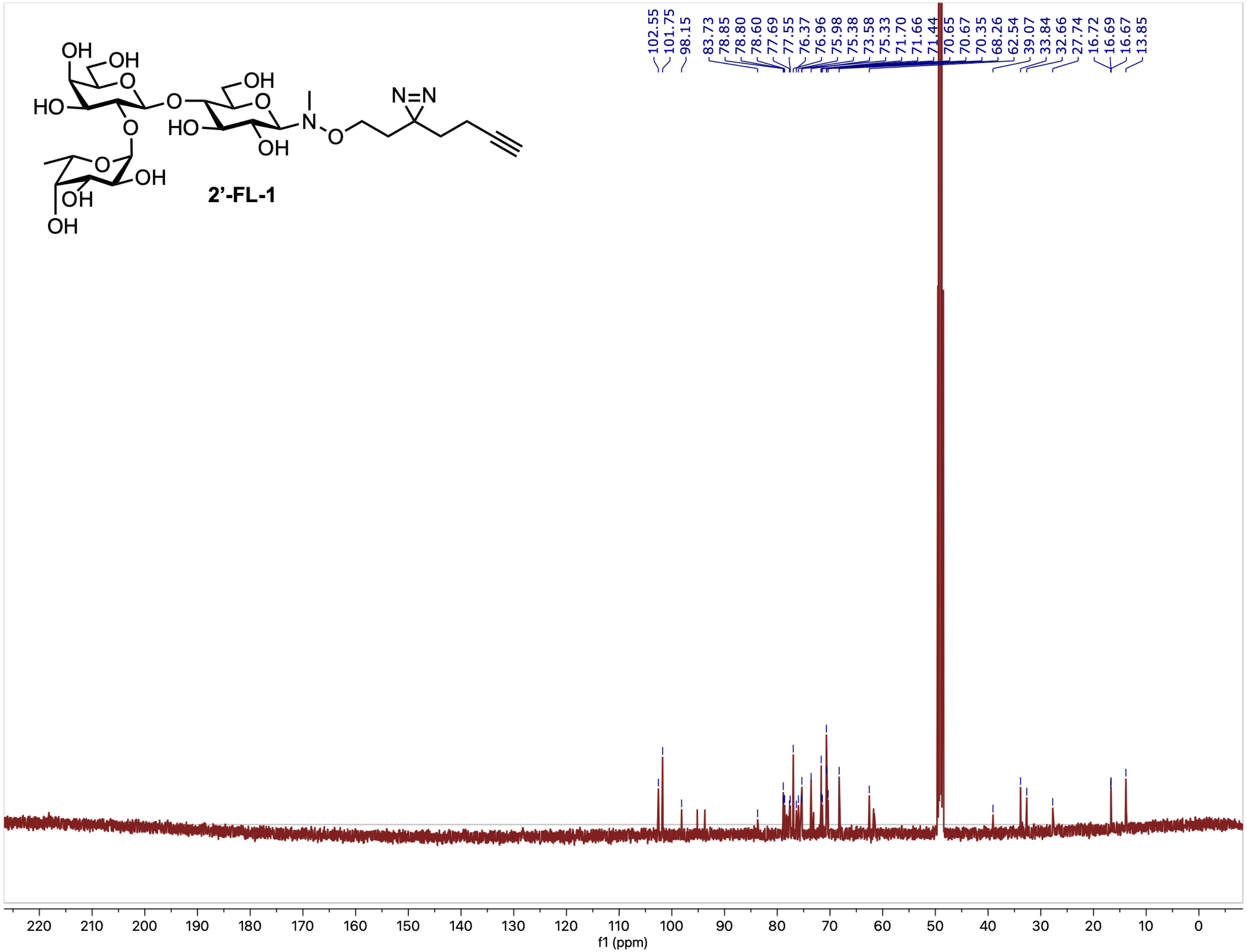

$^{1}H$ NMR (500 MHz, $CD_3OD$) spectrum of compound **3-FL-1**

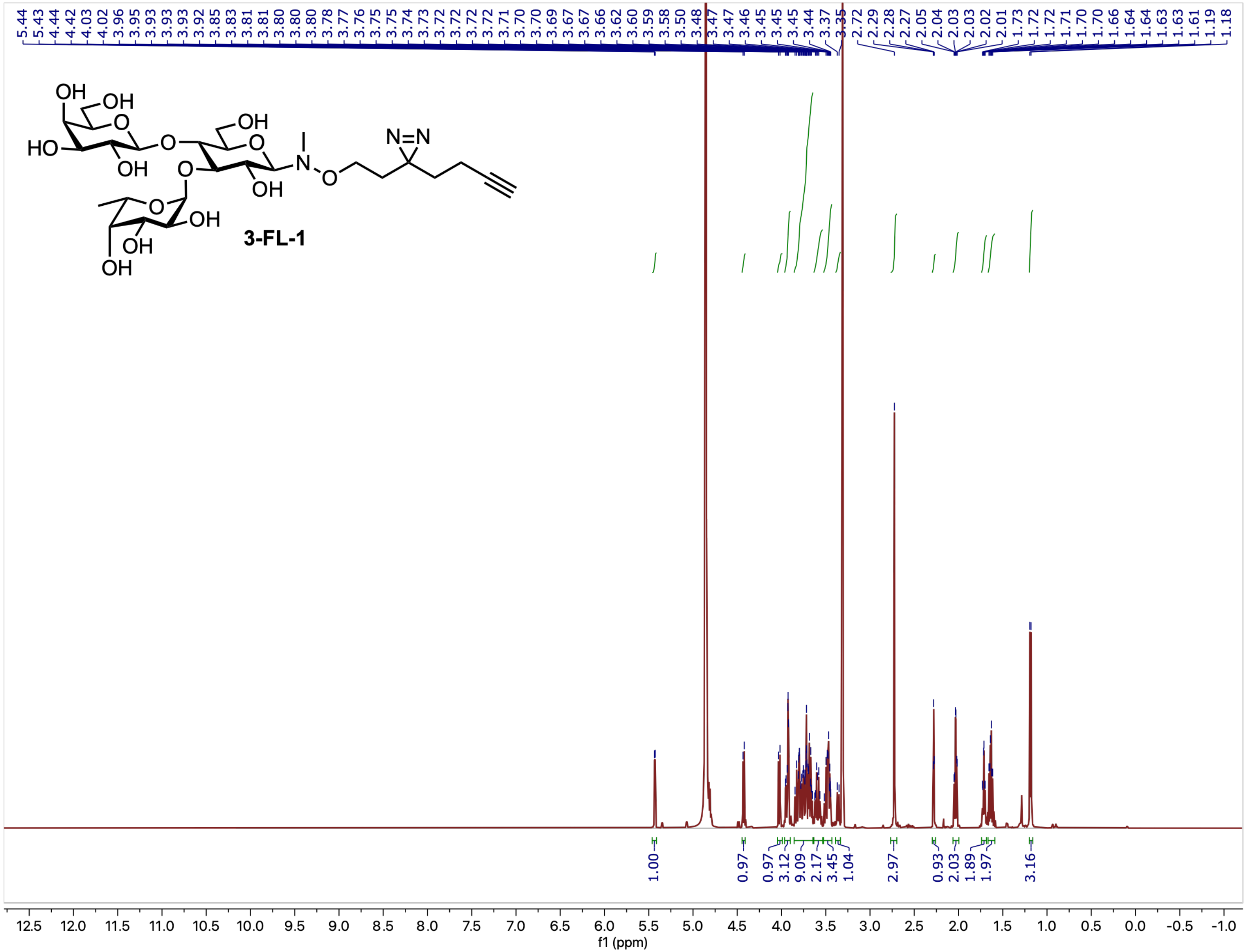

$^{3}$C NMR (126 MHz, $CD_3OD$) spectrum of compound **3-FL-1**

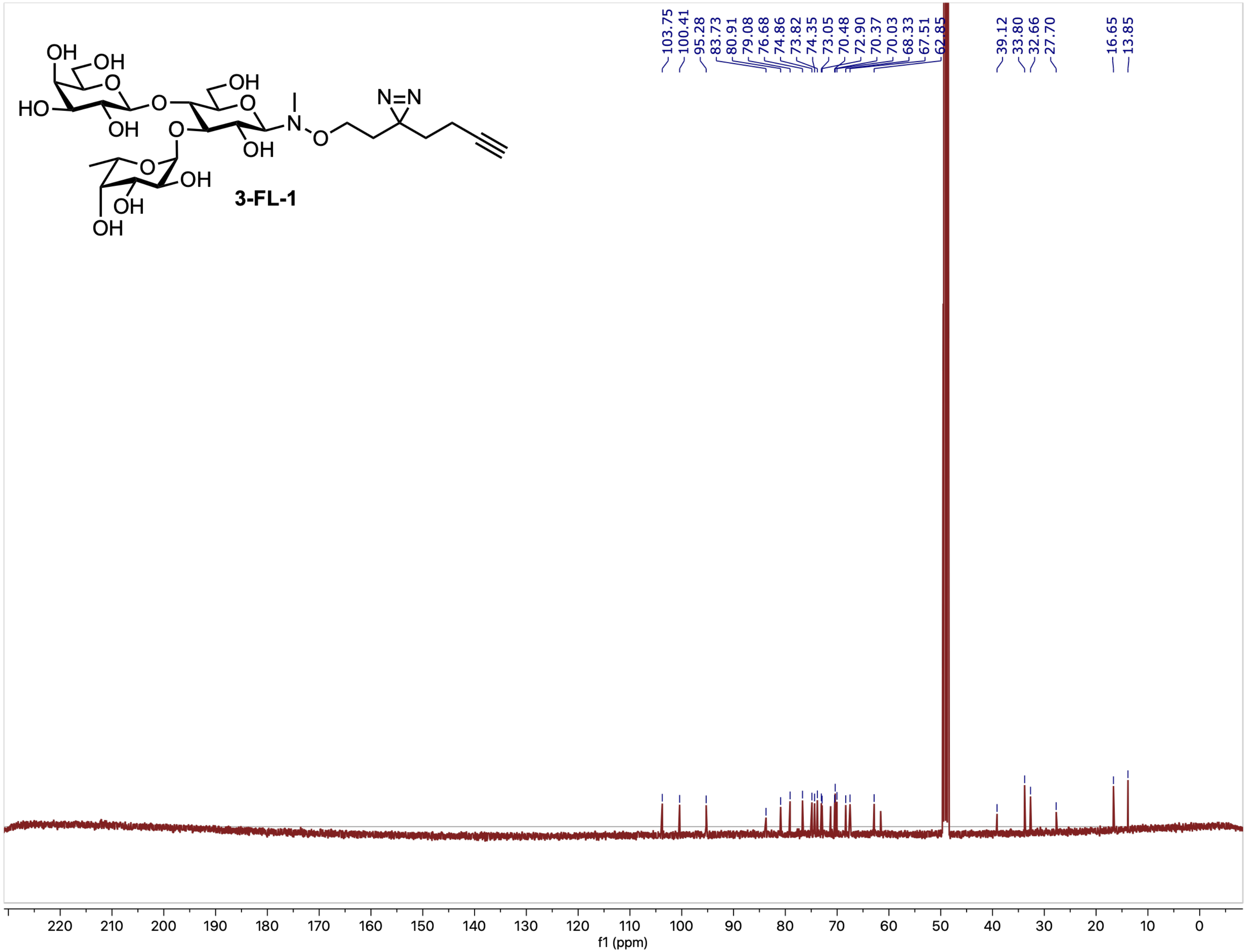
